# Task structure tailors the geometry of neural representations in human lateral prefrontal cortex

**DOI:** 10.1101/2024.03.06.583429

**Authors:** Apoorva Bhandari, Haley Keglovits, Defne Buyukyazgan, David Badre

**Author notes:** Corresponding author: Apoorva Bhandari. these authors contributed equally to this work (Apoorva Bhandari, Haley Keglovits). Competing Interests Statement: The authors declare no conflicts of interest.

## Abstract

How do human brains represent tasks of varying structure? The lateral prefrontal cortex (lPFC) flexibly represents task information. However, principles that shape lPFC representational geometry remain unsettled. We use fMRI and pattern analyses to reveal the structure of lPFC representational geometries as humans perform two distinct categorization tasks– one with flat, conjunctive categories and another with hierarchical, context-dependent categories. We show that lPFC encodes task-relevant information with task-tailored geometries of intermediate dimensionality. These geometries preferentially enhance the separability of task-relevant variables while encoding a subset in abstract form. Specifically, in the flat task, a global axis encodes response-relevant categories abstractly, while category-specific local geometries are high-dimensional. In the hierarchy task, a global axis abstractly encodes the higher-level context, while low-dimensional, context-specific local geometries compress irrelevant information and abstractly encode the relevant information. Trial-by-trial variability in neural patterns along the global coding axes in each task structure was associated with behavioral variability. Comparing these task geometries exposes generalizable principles by which lPFC tailors representations to different tasks.

## Introduction

Humans perform a diverse range of tasks in complex, variable settings. Performing each task requires attending to a varying set of relevant stimuli, actions, rules, goals, schedules and outcomes, organized by a cognitive strategy tailored to the rules and requirements of the task. How does the brain represent this task information in a way that supports the broad range of tasks people can perform?

The range of our expressive behavior relies on *cognitive control processes*^1,2^ that leverage circuits in the prefrontal cortex (PFC) to orchestrate sensory, motor, cognitive and affective processing across different brain regions in service of currently relevant task goals^3–6^. Mechanistic models of cognitive control posit that PFC neural activity enables this orchestration by encoding a *control representation* that uses task-specific inputs (stimuli, goals, rules, etc.) ^7–10^ to influence information processing throughout the brain. The vast range of potential human tasks means the lPFC must be capable of encoding control representations that implement tasks with widely varying structures.

Studies in non-human primates have provided evidence concerning the content of PFC control representations. Specifically, the activity of neurons located around the principal sulcus in the lateral PFC (lPFC) is sensitive to an array of information about whatever task is being performed, including task rules, stimulus features, responses, rewards, and other latent variables^11–17^. This principle of diverse coding of task information has been corroborated in the human brain using fMRI, where various types of task-relevant information can be decoded from hemodynamic activity in lPFC^18–24^. Previous work has also established that neural activity in lPFC and other regions as well as voxel activity in fMRI is broadly sensitive to the task dimension itself, such that neurons fire differently based on which task is being performed^25–30^. Therefore, a broad consensus has emerged that lPFC control representations are broadly task-sensitive, and encode most, if not all, task-relevant features.

However, while previous studies have found task-sensitive neural activity in lPFC, the coding principles that shape the *geometry* of these representations, and how they support task-specific computational demands across a range of diverse tasks, remain unsettled^31–37^. Understanding these principles is crucial because the geometry of a representation shapes what computations can be efficiently carried out with the encoded information and thus constrains whether it can be flexibly used across tasks.

A useful computational lens on lPFC coding and representational geometry comes from considering the *dimensionality* of the representation, which controls a tradeoff between the *separability* afforded by a neural code and the capacity for abstract, *generalizable* coding^35,36^

At one extreme, maximally compressed low-dimensional geometries encode each independent input feature along orthogonal dimensions. These dimensions are abstract in that they will encode the same feature regardless of the other inputs being encoded and can thus support generalization across independent input features. However, they do so at the cost of poor separability. In particular, such geometries will fail to support the readout of any potential task dimension that depends on nonlinearly mixing the input features and are therefore not useful for most tasks.

At the other extreme, maximally expanded high-dimensional geometries, constructed by nonlinearly mixing of all input features, achieve high separability for all potential combinations of task inputs. Therefore, these geometries are highly expressive and able to support the readout of any potential task dimension, but at the cost of generalizability. In particular, such geometries would be highly sensitive to small changes in the inputs, whether driven by noise or other task-irrelevant features.

Representational geometries between these two extremes provide a suite of possible compromises between the overall separability and generalizability of the representation. In particular, this intermediate regime includes geometries that would privilege both the separability and generalizability of some essential dimensions at the cost of other dimensions, thus affording representations that are adapted to the needs of the task^31^.

Considering this tradeoff between separability and generalizability, two contrasting accounts of coding principles of lPFC representations suggest different solutions to the problem of accommodating different task structures.

One account proposes that the lPFC learns specialized representations suited to each task encountered, storing a large repertoire of learned representations in the process^3^. To achieve this lPFC encodes the mix of inputs reflecting different task states on neural manifolds that are shaped, through learning, to optimize task-specific computations and the readout of task-relevant outputs. ^32,37–39^ (Fig.1). Such *task-tailored* representational geometries are shaped by the task’s computational demands, and can therefore span the entire range from low to high-dimensionality. Crucially, task-tailored representations selectively enhance the separability of task-relevant output dimensions, making them easier to read out. At the same time, they reduce separability along task-irrelevant dimensions to make the readout of task-relevant outputs robust to noise and generalizable over irrelevant features of the input^32,35,40,41^. Therefore, task-tailored representations combine the benefits of low and high dimensional geometries. However, these representations are specialized to the task and would not be suited to other tasks with differently structured input-output contingencies. Therefore, to adapt to different tasks, the population must learn to encode and read out from a variety of task-tailored representations and prevent interference between them^32,38^.

**Figure 1.**
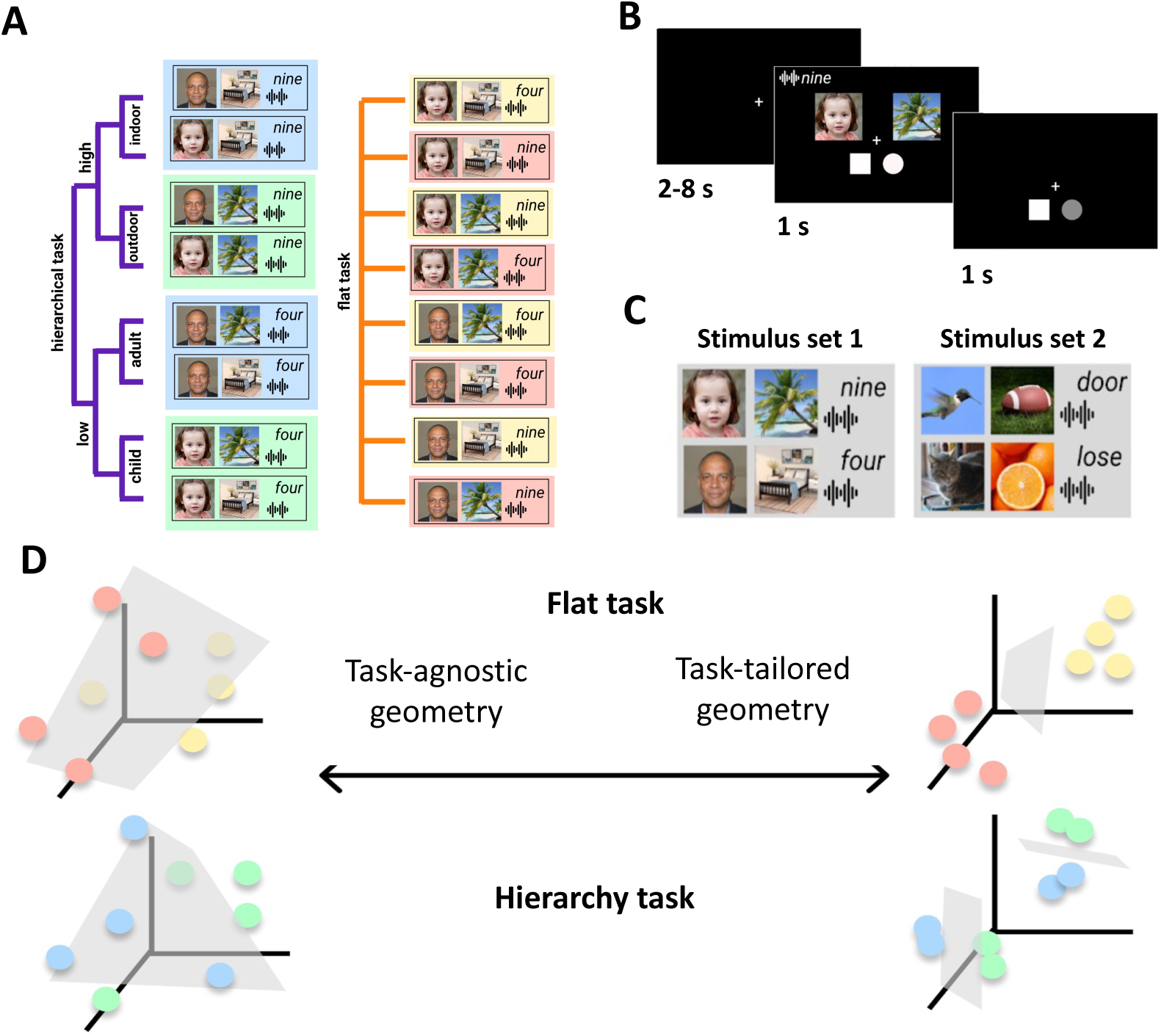
**(A)** Experimental tasks. Participants learnt two tasks with distinct structures – a hierarchy task (left) and a flat task (right). In the hierarchy task, one input feature specifies which of the other input features determines the response category hierarchically. In the flat task, the conjunction of all three input features determines the response category. **(B)** An example trial. The stimulus consists of a child’s face, an outdoor scene and the spoken number nine. The response panel consists of a square and a circle. If the subject selected the circle category, they would press the right key to indicate their response as the circle is on the right of fixation. **(C)** Stimulus categories employed. Different stimulus sets were employed for the two tasks for each participant and the stimulus–task mapping was counterbalanced across participants. **(D)** Task agnostic and task tailored representation hypotheses make distinct predictions for how the lPFC may accommodate tasks with different structures. Each plot represents a hypothetical neural space defined by the firing rates of 3 different neurons. The activity of the population in response to each trial types are shown as individual dots. Illustration of geometries for the flat task (top panel) and hierarchy task (bottom panel) under the two hypotheses. A task agnostic representational geometry employs a high-dimensional format which enhances separability across many different dimensions, allowing for multiple different readouts. Therefore, the same geometry can serve both tasks (left). Task-tailored representations leverage specifically enhance the separability of dimensions relevant to the task, while reducing it for irrelevant dimensions to enhance generalizability, such that the manifold is optimized for the task structure. Therefore, distinct geometries are learnt for each task.

An alternate account proposes that the lPFC employs an ‘adaptive’ coding scheme that rapidly establishes general-purpose representations that can be used to implement a wide range of tasks, regardless of their structure^34,42^. To achieve this, lPFC randomly mixes and projects its inputs onto high-dimensional manifolds that efficiently maximize the separability of all possible task structures arising from those inputs^34,35^. The higher separability enables a variety of arbitrary input-output contingencies to be read out from a single, expressive *task-agnostic representation*. This provides a basis for implementing multiple different input-output contingencies simultaneously^34^, with learning only needing to shape the readout, rather than shaping the representation. However, while such representations are expressive, they are also highly sensitive to all the features of the input, including irrelevant ones. This makes them less robust to noise, and less capable of generalizing to novel inputs.

While past studies have found extensive evidence for task-sensitivity of lPFC neural activity, the question of whether the geometries encoded in the activity are task-tailored or task-agnostic remains an open one. Past studies have found evidence for both low-dimensional and high-dimensional lPFC representations across different tasks, training regimes and species^31,32,34,35,43^. It has also been proposed that the lPFC can switch between low and high-dimensional representations depending on task demands^37^ or as the result of learning^40^. However, the mere observation of high dimensional geometry in lPFC does not provide evidence that its enhanced expressivity is used by the brain to accommodate multiple task structures, as opposed to other adaptive benefits of this geometry^36^. And likewise, the presence of low-dimensional geometry tailored to an individual task does not mean that new tasks are accommodated with separate low-dimensional manifolds. Indeed, no previous work has examined how lPFC representational geometry accommodates multiple tasks with different structures. Consequently, it remains unknown whether lPFC accommodates multiple tasks with an expressive, but task-agnostic representational geometry or task-tailored geometries specialized for individual task demands.

Crucially, while the task-tailored and task-agnostic representation accounts both predict task-sensitivity, they make different predictions about the geometry of representations across different tasks. Specifically, task-tailored representations should be specialized, and thus their geometries should differ between tasks of different structures. For each task, enhanced separability is predicted specifically for dimensions encoding task variables relevant to that task. On the other hand, task-agnostic geometries that can be shared across many task structures must employ higher-dimensional formats that enhance the separability of multiple dimensions simultaneously, at the cost of generalizability^35^. The same task-agnostic representation would be used for different tasks.

Here we test the predictions of these two accounts by examining how lPFC neural representations adapt to the demands of two tasks with structurally different input-output contingencies. We extensively trained human participants to perform two distinct categorization tasks (Fig. 1). One was a flat task where the categorization was defined based on a non-linear mapping. The other was a hierarchical task where participants switched between two stimulus-defined contexts, with the categorization defined by independent stimulus features across contexts. Using decoding and representational similarity analysis of fMRI data, we characterized the lPFC representational geometries of these tasks at a level of detail unprecedented in humans. We found evidence that lPFC encodes task-tailored, rather than task-agnostic geometries, which were shaped by task structure based on principles that balance separability and generalizability.

## Results

### Experimental tasks and training

To investigate representational geometries in lPFC, we trained human participants (N = 94; 58 female, 33 male, 3 declined to answer; mean age = 22.9 ± 4.7) on two categorization tasks with different structures. In each task, a set of eight types of stimuli defined by two visual features (e.g., adult or child face, indoor or outdoor scene) and an auditory feature (e.g., low or high spoken number words), were mapped to two response categories symbolized by visual shapes e.g., circle or square). On presentation of the stimulus, participants had to decide to which category the stimulus belonged and indicate their choice by pressing a key based on the location of the associated category symbol. Participants were first trained on a ‘flat’ task in which categorization required a consideration of the conjunction of all three stimulus features (specified by a latent XOR rule, see Methods), all of which were necessary for identifying the category (Fig 1). They were then trained on a ‘hierarchy’ task with a separate set of stimuli in which the mapping was defined by a hierarchical rule, such that the auditory feature dictated which of the two visual features was relevant for identifying the category (Fig 1).

Note that the two tasks were assigned different stimuli in each participant and the assignment of stimulus set to task was counterbalanced across participants. For convenience, we will henceforth refer to the three stimulus features as visual feature 1 (young/old face or mammal/bird), visual feature 2 (indoor/outdoor scene or edible/inedible object) and auditory feature (low/high number or noun/verb). We will refer to the category (square/circle or triangle/pentagon) which participants used to select their response as the response category, and to the physical button press (index or middle finger) as the motor response.

Additionally, our stimulus sets were designed to concurrently and systematically vary along three orthogonal task-irrelevant dimensions. For example, while the task-relevant dimension for objects was edible vs inedible, the objects also varied along an orthogonal natural vs man-made dimension. Similarly, while low/high magnitude was a task-relevant dimension, the auditory stimuli were also odd or even (see Methods for all orthogonal dimensions). These orthogonal, task-irrelevant features (two visual, 1 auditory per stimulus set) were never relevant for performing the task and were never described to participants. These orthogonal dimensions were included in order to test the effect of task-relevance on representational geometries.

Participants were trained on the tasks until they reached a minimum overall accuracy of 85%, as well as an accuracy of 80% on each of the eight individual trial types. During this training phase, participants were both explicitly instructed on the stimulus-category mappings, as well as given trial-by-trial feedback. Participants took more trials on average to achieve the predefined criterion on the flat compared to the hierarchy task (flat task: mean 1416, range 440-2640 trials-to-criterion; hierarchy task: mean 409 trials, range 176-1056 trials-to-criterion).

A subset of the participants that met pre-defined performance criteria on both tasks then performed approximately 2000 trials for each task over 10 days (5 days for each task), while being scanned with fMRI (N = 20, 14 female, 6 male, mean age = 22.8 ± 4.6; see Methods for full exclusion criteria). During the training phase, the scanned participants exhibited terminal error rates that were significantly higher for the flat task than the hierarchy task (flat task error rate: 9.5%, hierarchy task error rate: 6.4%, paired t-test: *t =* 3.87, *p* = 0.001), as well as terminal response times (flat task mean RT 1398 ms, hierarchy task mean RT 1277 ms; paired *t*-test: *t =* 4.25, *p* < 0.001) (Fig. 2), which was expected as the hierarchical rule simplifies the response selection problem.

**Figure 2.**
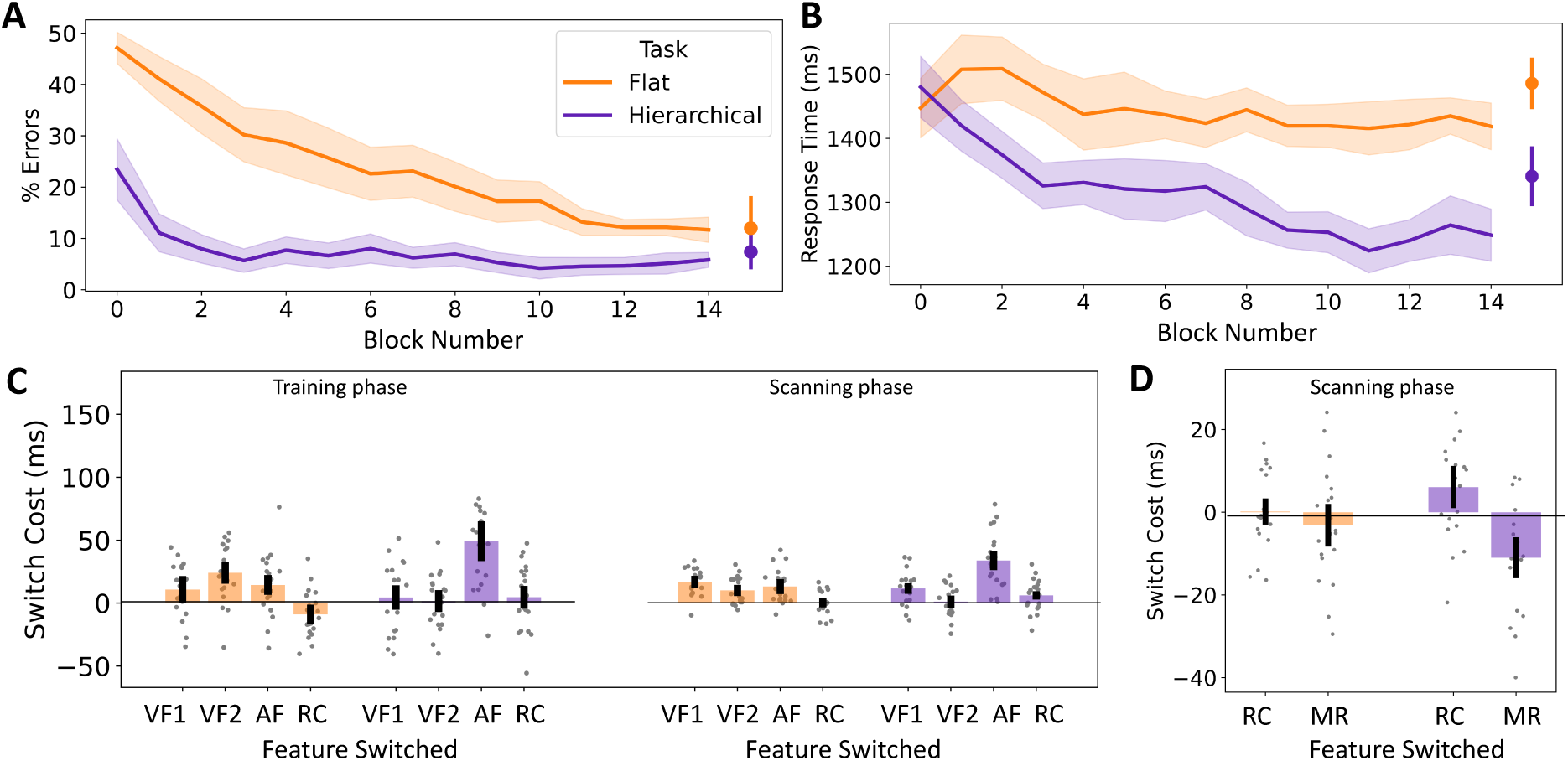
Behavioral performance of scanning group. Error rates **(A)** and response times **(B)** during training (lines) and scanning (dots) phase. Participants were slower to learn and had a higher asymptotic error rate on the flat task at the end of the training phase. Error rates increased slightly in the scanner phase. Lines represent block-wise performance in the training phase. Dots represent mean overall performance in the scanning phase. Participants did not show an improvement in response time on the flat task. Participants were significantly slower to respond in the scanner (points on the right) compared to the end of training. **(C)** Behavioral evidence that participants used distinct strategies in flat vs hierarchy tasks. During training (left plots), in the flat task, RT costs were similar for switching of the different features. In the hierarchy task, there was a significantly larger cost when the auditory dimension (AF), which signaled superordinate context, switched. This pattern of hierarchical switching across tasks was maintained during the scanning phase (right plots). **(D)** In the scanning phase, response category and motor response interacted in driving RT. VF1 = Visual Feature 1, VF2 = Visual Feature 2, AF = Auditory Feature, RC = Response Category, MR = Motor Response. All error bars reflect 95% confidence intervals.

### Behavioral performance on the flat and hierarchy task

Following training, participants continued to perform at a high accuracy on both tasks in the subsequent scanning sessions, despite no longer receiving trial-by-trial feedback (Fig. 2; flat task error rate: 14%; hierarchy task error rate: 8%). Performance was slower overall in the scanner (mean flat RT 1468 ms, mean hierarchy task RT 1335 ms), as is commonly observed^44,45^. A repeated measures ANOVA of accuracy confirmed a main effect of task (*F_1,57_* = 22.5, *p* < 0.001 and phase (*F_1,57_* = 11.2, p = 0.001) but no task:phase interaction (*F_1,57_* = 1.6). A similar pattern was observed for RT with significant main effects of task (*F_1,57_* = 52.3, *p* < 0.001 and phase (*F_1,57_* = 14.9, p < 0.001) but no task:phase interaction (*F_1,57_* = 0.6).

We tested whether participants employed distinct strategies across the two tasks by analyzing the pattern of RT costs associated with switches of different task features (Fig. 2C). In a task-specific ANOVA on the flat task, while switch costs were evident when a feature switched (main effect of switching: *F_1,114_ =* 117.2*, p <* 0.001), those costs did not significantly vary across the features (switch:feature interaction: *F_2,114_ =2.17*). In contrast in the hierarchy task, we observed a significantly larger cost of switching the auditory feature which signaled a switch in the higher-level context than either of the visual features (switch:feature interaction: *F_2,114_* = 25.9, *p* < 0.001), confirming that participants employed the instructed hierarchical strategy for solving the task despite extensive practice. These differences in switching effects across tasks were supported by a significant three-way task (flat vs hierarchy):trial-type (switch vs repeat):feature (stimulus dimensions 1-3 or response categories) interaction (*F_5,228_* = 7.5*, p <* 0.001) along with significant main effects of task (*F_1,228_* = 38.3*, p <* 0.001), switching (*F_1,228_ =* 117.2, *p <* 0.001) and the switch:feature interaction (*F_2,228_ =* 33.5*, p <* 0.001).

We did not observe a significant effect of switching the response category in either task (*p* > 0.1). Further analysis revealed a significant interaction between response category switching and motor response switching (Fig. 2D) in both the flat (*F_1,57_* = 23.6, *p* < 0.001) and hierarchy (*F_1,57_* = 49.8, *p* < 0.001) tasks. In both tasks, the lack of an observed effect of switching the response category was driven by a pattern of partial overlap costs^46^, such that switching any one of the features produces a large cost, but switching both features or no features produced lower costs.

### Left lPFC encodes diverse task-relevant information

We preselected a cluster of 10 parcels in the mid and dorsal portion of left and right lPFC from a previously published cortical parcellation^47^. These parcels cluster as part of a frontoparietal network defined by resting state connectivity^48^ and overlap with loci previously associated with cognitive control^5^. Moreover, their resting state functional connectivity pattern resembles that of lPFC regions studied in macaques^49^ showing robust evidence for the diverse, high dimensional coding of task information^34^, which provides the premise for our hypotheses.

Given the difficulty of decoding from lPFC with human fMRI^50^, we maximized power in our analyses by pooling all the voxels across the 5 lPFC parcels in each hemisphere to create merged left lPFC and right lPFC ROIs. These parcels all belong to the same resting state network and thus have correlated fluctuations in activity and our hypotheses do not predict qualitative differences across them. An alternate approach where results were averaged across parcels within each hemisphere produced similar results (Supplementary section S3). In addition to the left and right lPFC ROIs, we also selected parcels associated with the left and right primary auditory cortex (pAC). This region is involved in early sensory processing, serving as a control for lPFC.

Whole-brain, voxel-wise general linear models (GLMs) were employed to estimate the hemodynamic response associated with the 8 trial types in each scanner run, separately for both tasks. GLMs were conservatively designed to minimize bias in the training set and control for confounding factors like motor responses and the number of trials that went into estimating the run-wise responses to each trial type (see Methods). From the resulting beta images, ROI-specific multi-voxel patterns were extracted and analyzed using both decoding and representational similarity analysis approaches.

We first examined the content of neural representations in lPFC and pAC for comparison. Under the task-agnostic geometry hypothesis, all task-relevant information, and potentially task-irrelevant information, is predicted to be decodable in lPFC and the information content of the representations should not vary across the two task structures. Under the task-tailored hypothesis, differing task demands imposed by the task structures would shape the content of the representations resulting in qualitative or quantitative differences in the information decodable across the two task structures.

We employed cross-validated multivoxel decoding to test whether local patterns in the lPFC parcels encode task-relevant information. Using a leave-one-run-out cross-validation scheme, linear support vector machines (SVMs) were trained to decode task-relevant dichotomies from ROI-specific multi-voxel patterns associated with the 8 trial types. For each task, we trained decoders for each of the three stimulus features, the response category and the motor response. In each cross-validation fold, the decoders were tested on the patterns from the left-out run and decoding accuracies were averaged across cross-validation folds. Training data were always carefully balanced to avoid any classifier biases (see Methods). Classifier performance was evaluated against chance (50%) with permutation tests and compared across conditions with parametric statistical tests.

In line with previous results, we found evidence for the coding of diverse task-relevant information in lPFC (Fig 3A,C). As expected for fMRI decoding in lPFC^50^, mean decoding accuracies in left lPFC ROI were low but significantly above chance levels (permutation tests, correcting for multiple comparisons for the number of ROIs tested– i.e. for 4 ROIs) for most task features across both the flat task (visual feature 1: 51.9%*, p* = 0.005*; visual feature 2: 56.2%, *p* < 0.001*; auditory feature: 53.1%, *p* < 0.001*; response category: 53.9%*, p* < 0.001*; motor response: 50.1%, *p* = 0.43) and the hierarchy task (visual feature 1: 51.8%, *p* = 0.003*; visual feature 2: 52.7%, %, *p* < 0.001*; auditory feature/context: 61.5%, *p* < 0.001*; response category: 54.4%, *p* < 0.001*; motor response: 51.77%, *p* = 0.004). In the right lPFC, all task features barring the auditory feature were decoded above chance levels in the flat task (visual feature 1: 52.5%, *p* < 0.001*; visual feature 2: 51.6%, *p* = 0.007*; auditory feature/context: 61.5%, *p* < 0.001*; response category: 52.5%, p < 0.001*; motor response: 50.53%, *p* = 0.21) while in the hierarchy task, the auditory feature (reflecting context), visual feature 2 and the response category were decoded above chance levels (visual feature 1: 50.9%, *p* = 0.151; visual feature 2: 51.4%, p = 0.013; auditory feature/context: 58.9%, p < 0.001*; response category: 52.2%, p < 0.001*; motor response: 50.9%, *p* = 0.1).

**Figure 3.**
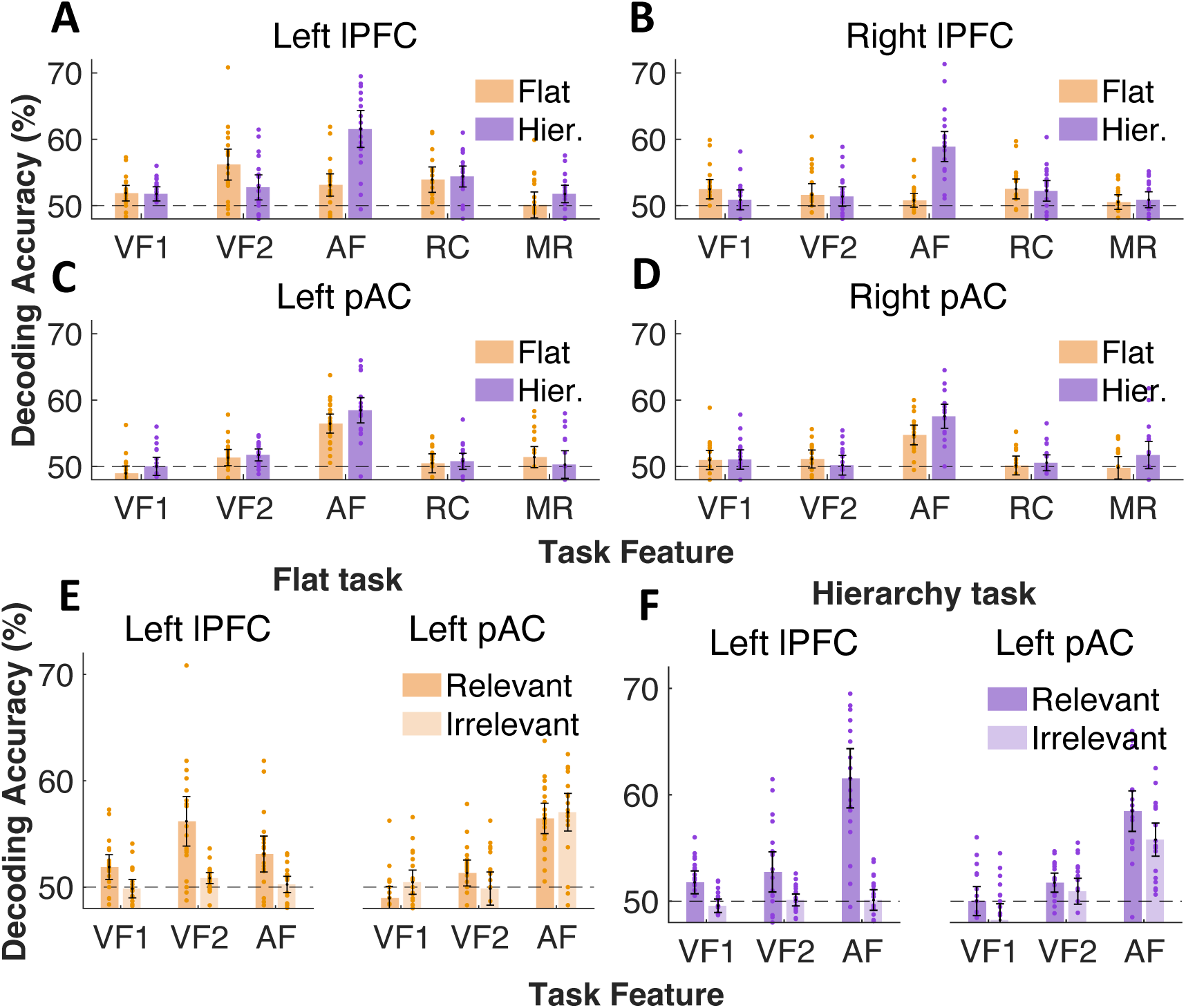
Information content of lPFC and pAC representations. Cross-validated decoding accuracies for all task features in left lPFC (A), right lPFC (B), left pAC (C) and right pAC (D). While left lPFC shows diverse coding of various task features, left and right pAC only show coding of auditory stimulus information. Across both, the flat task (E) and hierarchy task (F), left lPFC shows selective coding of task-relevant stimulus information. On the other hand, left pAC shows obligatory coding of auditory stimulus information regardless of its relevant for the task. VF1 = Visual feature 1, VF2 = Visual Feature 2, AF = Auditory Feature, RC = Response Category, MR = Motor Response. All error bars reflect 95% confidence intervals.

Task-specific mixed model ANOVAs comparing decoding accuracies in left lPFC across sub-groups of participants that were assigned different stimulus sets for each task showed that the diverse coding of task-relevant features was insensitive to this assignment in both the flat (*F*_1,18_ = 0.2) and hierarchy task (*F*_1,18_ = 1.2).

A repeated measures ANOVA of the left lPFC decoding accuracies with task and feature as factors revealed a significant task:feature interaction (*F*_4,76_ = 11.9, *p* < 0.001). Posthoc comparisons found significantly higher decoding accuracies specifically for the auditory feature in the hierarchy task compared to the flat task (Bonferroni corrected *p <* 0.001). Therefore, while task structure did not appreciably influence what information was coded in the lPFC, it did influence the strength of coding of the auditory feature which plays the role of superordinate context in the hierarchy task, but not the flat task, an observation consistent with the task-tailored representation hypothesis.

In contrast to the lPFC, we did not observe diverse coding in the primary auditory cortex (pAC, Fig. 3B,D). Only the auditory stimulus feature was strongly coded in the pAC across both the left (flat task 56.5%, *p <* 0.001*; hierarchy task: 58.5%, *p <* 0.001*) and right hemispheres (flat task: 54.8%, *p <* 0.001*, hierarchy task: 57.6%, *p <* 0.001*). Similar to the lPFC, however, the auditory feature in pAC was also shaped by task structure, with the accuracies being higher specifically in the hierarchy task (where the auditory feature signals context) compared to the flat task (*F*_1,19_ =11.6, *p* = 0.003).

In summary, these results replicate previous reports of the coding of diverse, task-relevant information in lPFC (particularly in the left lPFC). These results were specific to the left lPFC and markedly differed from the primary auditory cortex, which preferentially coded auditory stimulus information. Moreover, while task structure did not influence what information was decodable it strongly influenced the strength of decoding of the auditory feature in the hierarchy task. While this difference is consistent with the task-tailored representation hypothesis, it is also consistent with greater attention or processing time spent on this feature, which likely resulted in the same pattern in pAC. Subsequent analyses were designed to provide more specificity.

### Left lPFC preferentially encodes task-relevant information across the flat and hierarchy task

We next asked whether lPFC preferentially codes task-relevant information. An extreme version of the task-agnostic representation hypothesis is that lPFC obligatorily receives and non-linearly mixes a wide variety of task-relevant and task-irrelevant information^51^. Such a representation would reflect a coding strategy that does not commit to any particular task or task structure, thus affording maximum flexibility in adapting to any task at hand. Alternatively, the task-tailored representation hypothesis predicts a strong preference for coding task-relevant information.

To test whether lPFC preferentially codes task-relevant information, we leveraged the fact that all stimuli used in the experiment were also systematically varied along orthogonal dimensions that were not used in the task or discussed with participants. For example, the task-relevant feature for objects was edible vs inedible, while the orthogonal irrelevant feature was animate vs inanimate. It is important to emphasize that, by design, these orthogonal dimensions were *never* task-relevant. Results are plotted in Fig. 3E and 3F.

In the left lPFC, we found little evidence for the coding of orthogonal task-irrelevant stimulus features with no mean decoding accuracy being reliably above chance levels, except for visual feature 2 in the hierarchy task (visual feature 2: 51.8%, *p* = 0.011*) which was driven by the irrelevant feature of scene stimuli with or without people (53.2%, *p =* 0.002*). Task-specific, stimulus feature x task relevance rmANOVAs confirmed a main effect of task relevance in both the flat task (*F*_1,19_ =37.2, *p* < 0.001) and the hierarchy task (*F*_1,19_ =71.6, *p* < 0.001). In the hierarchy task, the effect of task relevance interacted with stimulus feature (*F_2,38_* = 18.3, *p* < 0.001), with Bonferroni corrected posthoc tests showing a stronger effect for the auditory feature (*p* < 0.001) and visual feature 1 (*p* = 0.002) and visual feature 2 (*p* = 0.026), which was driven by lPFC coding of the irrelevant natural vs manmade category.

In stark contrast with the lPFC, pAC coded both task-relevant and orthogonal, irrelevant auditory stimulus features. We found strong coding of the orthogonal, task-irrelevant auditory features in both left (flat task: 57.0%, *p <* 0.001*; hierarchy task: 55.8%, *p <* 0.001*) and right (flat task: 53.9%, *p <* 0.001*; hierarchy task: 54.3%, *p <* 0.001*) primary auditory cortex. Therefore, unlike the lPFC, pAC coded both task-relevant and task-irrelevant auditory stimulus features. Similar to the lPFC, however, the strength of coding of auditory features in pAC was shaped by task structure. Specifically in the hierarchy task, decoding accuracies for the task-relevant auditory feature were reliably higher than those for the orthogonal, irrelevant auditory feature (*F*_1,19_ =13.3, *p* = 0.002). In the flat task, we found no such evidence for an effect of task-relevance (*F*_1,19_ =0.03, *p* > 0.1).

Collectively, these results support the conclusion that left lPFC preferentially encodes task-relevant information. On the other hand, the pAC only encoded auditory information, regardless of whether it is task-relevant.

### Left lPFC representations show enhanced separability of task-relevant variables across task structures

The task-tailored and task-agnostic hypotheses make distinct predictions about the separability of lPFC representations. While the task-agnostic hypothesis predicts high separability along all dimensions, the task-tailored hypothesis predicts enhanced separability of the key dimensions coding task-relevant variables. Importantly, the task-tailored hypothesis predicts that while overall separability may be similar across both tasks, different task variables will show high separability across the two tasks, owing to their distinct structure. To test these hypotheses, we focus specifically on the left lPFC where we observed diverse coding of task-relevant information, and for comparison, on bilateral pAC, where we observed preferential coding of auditory information. Given that left and right pAC showed similar representational content, we averaged across them in subsequent analyses.

To assess the separability of lPFC representations, we examined decodability of all possible binary dichotomies with a linear classifier. These binary dichotomies reflect the many possible ‘dimensions’ that could be read out from the mixture of the task inputs. The greater the number of such dichotomies that can be read out, the more separable the representation^34,35^. To minimize classifier bias resulting from unbalanced training data, we restricted ourselves to all 35 balanced binary dichotomies which had an equal number of patterns for each class (4 trial types each). ^31,40^ A maximally compressed, low-dimensional representation consisting of only pure selective neurons would support the coding of fewer than 5 dichotomies^35^ while a maximally expanded, high-dimensional representation would support all 35.

The decodability of dichotomies was evaluated at the group level with permutation tests. After Bonferroni correction for multiple comparisons for the number of dichotomies, 27/35 dichotomies in the flat task and 21/35 dichotomies in the hierarchy task were decodable above chance levels in the left lPFC at the group level with no significant difference across the two tasks (*t* = 1.5, p = 0.14). In contrast, a parallel analysis of the representation of the three orthogonal, task-irrelevant features found that, after Bonferroni correction for multiple comparisons, none of the 35 dichotomies formed from only the task-irrelevant features could be decoded above chance levels for either of the tasks. Even at a more liberal threshold (i.e., without multiple comparison correction), 0/35 (flat task) and 2/35 (hierarchy task) dichotomies could be decoded. Therefore, the observed proportions find evidence that only the task-relevant inputs across both task structures were encoded with geometries of intermediate dimensionality.

We also compared the separability across the two tasks using *shattering dimensionality,* defined as the average decoding accuracy across all 35 dichotomies. Shattering dimensionality reflects the overall separability of the representation and is informed by separability along a wide range of neural dimensions. Shattering dimensionality was reliably above chance (Fig 4B) for both the flat task (52.8%, *p* < 0.001*) and the hierarchy task (52.6%, *p* < 0.001*). Again, shattering dimensionality did not reliably differ across the two tasks (paired *t*-test: *t =* 0.54), with Bayes factor offering moderate evidence in favor of the null hypothesis (BF_10_ = 0.26).

**Figure 4.**
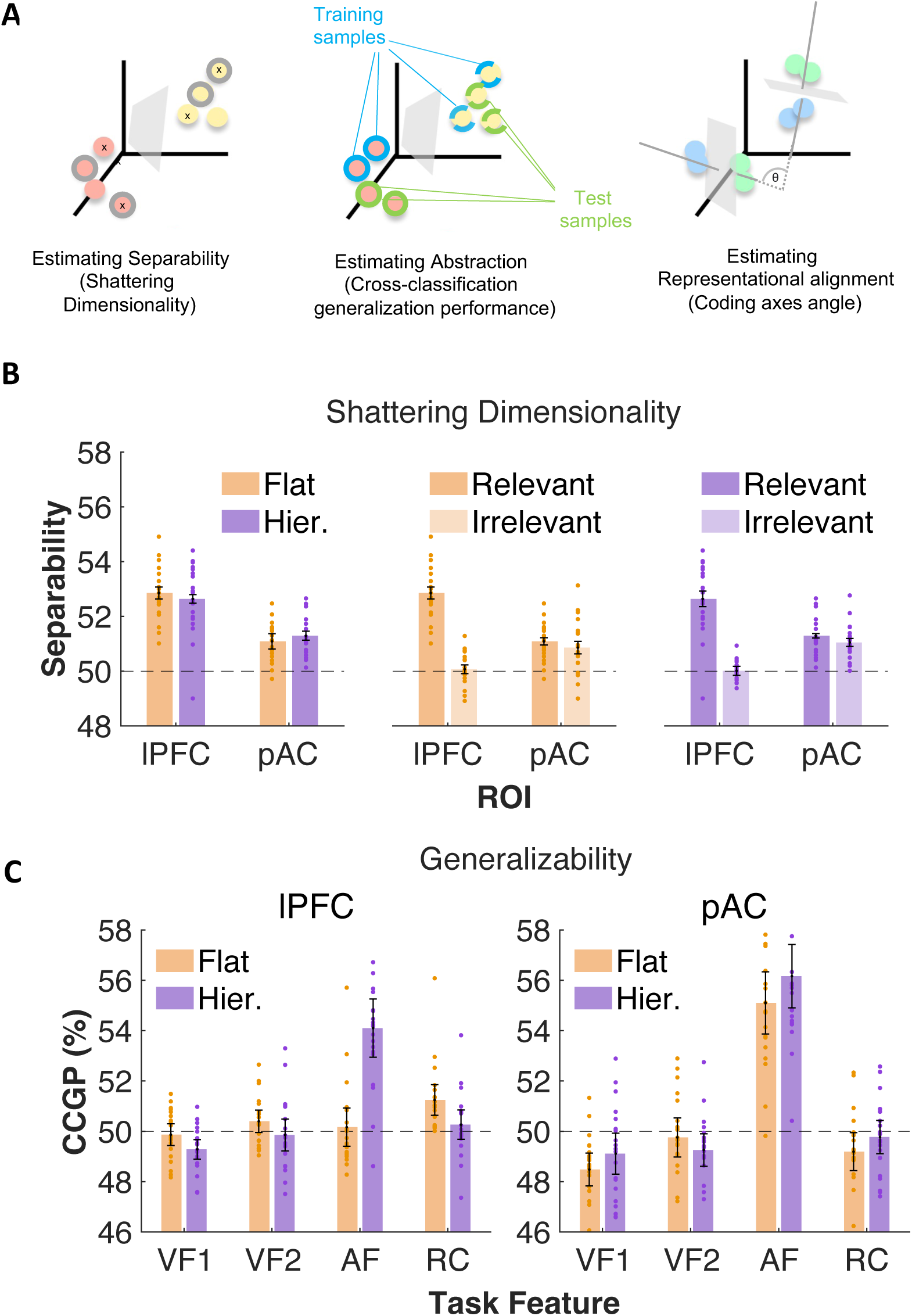
**(A)** Separability (left panel) was assessed by decoding all possible balanced dichotomies. The same set of 8 patterns can be split into different dichotomies or classes. For example, three dichotomies are illustrated as either red vs yellow points, outlined vs not outlined points, and points marked with x vs unmarked. A total of 35 balanced dichotomies are possible. Shattering dimensionality is the mean cross-validated decoding accuracy averaged across all dichotomies. Generalizability or abstraction was assessed through cross-generalization (middle panel). Classifiers were trained on half the points assigned to each label (blue in middle panel), and tested on the other half (green in middle panel), with all possible combinations of training and test sets evaluated. The averaged cross-classification generalization performance (CCGP). Finally, cross-cluster representational alignment is assessed by measuring the angle (θ) between the coding axis associated with the same feature in the two clusters. The coding axis is mathematically the weight vector of the trained classifier. **(B)** Shattering dimensionality, defined here as the mean cross-validated decoding accuracy across all 35 balanced binary dichotomies, is a measure of the separability of the representation. The separability of representations in lPFC and pAC was significantly above chance levels, and was not different across tasks or regions (B, left panel). Separability for the representation of the task-relevant inputs was significantly higher than for the orthogonal, task-irrelevant inputs in both flat (B, middle panel) and hierarchy (B, right panel) tasks. **(C)** Different task variables were encoded in an abstract, generalizable form in the lPFC (C, left panel) across the two tasks. In the flat task, the response category (RC) was encoded in abstract form, while in the hierarchy task it was the auditory feature (AF). In the pAC (C, right panel) only the auditory feature was abstractly coded. VF1 = Visual feature 1, VF2 = Visual Feature 2, AF = Auditory Feature, RC = Response Category. All error bars reflect 95% confidence intervals.

While shattering dimensionality provides a measure of overall separability reflecting all neural dimensions, it can be biased by the strength of pure-selective coding of task variables. To examine separability while minimizing the influence of pure selective coding, we identified dichotomies in each task that were orthogonal to all four task-relevant variables (3 stimulus features and the response category). The separability of these dichotomies cannot be influenced by pure selective coding of any of the task variables. This *mixed-selective separability* was reliably above chance in the left lPFC only for the flat task (51.5%, *p* < 0.001*) and not the hierarchy task (50.2%, *p* = 0.34) providing some evidence for non-linear mixed selectivity in the flat task in lPFC. However, this measure also did not reliably differ between the tasks (paired *t*-test: *t =* 1.9, p = 0.08) with the Bayes Factor indicating equivocal evidence (BF_10_ = 0.99). Therefore, there was no clear evidence that the separability of the lPFC task representations is shaped by task structure.

Of the 35 dichotomies, 4 are identifiable task variables (three stimulus features, response category), while the rest reflected arbitrary mixtures of these variables. The task-tailored representation hypothesis predicts that separability should be enhanced specifically for dimensions encoding task variables while the task-agnostic representation hypothesis predicts equivalent separability across many dimensions, including those that may not be required to implement the task. Separability in left lPFC parcels was significantly higher for the dichotomies encoding the task variables compared to the 31 other arbitrary variables in both the flat task (paired *t*-test, *t =* 2.9, *p* = 0.009*) and the hierarchy task (paired *t*-test, *t =* 7.6, *p* < 0.001). Therefore, in left lPFC, separability is preferentially enhanced for neural dimensions encoding task variables.

In the pAC (averaged across the two hemispheres), like the lPFC, shattering dimensionality was again reliably above chance for both the flat task (51.1%, *p* < 0.001*) and the hierarchy task (51.3%, *t* = 7.8, *p* < 0.001*), and it did not differ between the two tasks (*t* = 1.2, BF_10_ = 0.42). Moreover, separability was higher for neural dimensions encoding task variables (flat task: *t* = 3.4, *p* = 0.003*; hierarchy task: *t* = 6.43, *p* < 0.001*). However, unlike left lPFC, paired *t-*tests showed no reliable difference in the separability of the task-relevant features vs the orthogonal, task-irrelevant features (flat task: *t* = 0.9, BF_10_ = 0.34; hierarchy task: *t* = 1.17, BF_10_ = 0.42) in pAC. Therefore, the overall separability of pAC representations is not shaped by either task-relevance or task structure.

We next tested the differences in representational geometry between lPFC and the pAC. Shattering dimensionality was not reliably different between the left lPFC and pAC (flat task: *t* = 0.69, BF_10_ = 0.29; hierarchy task: *t* = 1.3, BF_10_ = 0.46). However, this comparison is potentially biased by differences across regions in the signal-to-noise and the strength of pure selective coding. In particular, the shattering dimension might reflect strong pure selective coding of auditory information in the pAC, while in the lPFC the measure reflects non-linear mixed selective coding of a variety of task variables. Therefore, we examined mixed-selective separability which is not influenced by pure selective coding of any of the task variables. A region x task rmANOVA confirmed a significant main effect of region (*F*_1,19_ =15.2, *p* < 0.001) on mixed selective separability, with higher separability in the lPFC compared to the pAC and no main effect of task or interaction with task (*p* > 0.1).

Collectively, these results support the conclusion that left lPFC selectively encodes task-relevant information with a geometry of intermediate dimensionality, non-linearly mixing task-relevant information to enhance separability along several neural dimensions. In support of the task-tailored hypothesis, separability in the lPFC was preferentially enhanced for neural dimensions encoding task variables. On the other hand, the pAC employs a low-dimensional geometry with pure selective coding primarily of auditory information, regardless of whether it is task-relevant.

### Task structure shapes the generalizable coding of task features in lPFC

A key feature of task-tailored representations is supporting the generalization of certain task variables and their combinations at the cost of separability along other variables or their combinations. We next assessed the extent to which left lPFC representational geometries support generalization. We again trained linear SVMs to decode the three stimulus features and the response category using a leave-one-run-out cross-validation procedure. However, this time, the SVMs were trained on the patterns associated with half the trial types (two for each level of the decoded feature), and tested on the patterns associated with the remaining trial types from the left-out test run. Therefore, the SVMs were evaluated on their *cross-classification generalization performance* (CCGP, Fig 4A, right panel), a measure that assesses the extent to which the coding of the relevant feature is abstract^31^. For the maximally high-dimensional geometries where different task states are randomly distributed in multi-dimensional neural space, CCGP is expected to be at chance levels. For lower-dimensional geometries in which task states sharing the same feature tend to cluster together, CCGP is expected to be systematically above chance levels^31^.

The two tasks differed in terms of which task variables were abstractly coded by this measure (Fig 4C, left panel). In the flat task, CCGP was reliably above chance for the response category in left lPFC (53.5%, *p* < 0.001) and for visual feature 2 (52.1%, p < 0.001) but not for any of the other stimulus features (all *p*s > 0.1). In the hierarchy task, CCGP was reliably above chance levels in left lPFC for the auditory/context feature (left lPFC: 60.3%, *p* < 0.001), and the response category (51.4%, p < 0.001) but not for any of the stimulus categories. Paired *t*-tests confirmed that CCGP for the auditory dimension was significantly higher in the hierarchy task than the flat task (*t =* 5.8*, p* < 0.001) while CCGP for the response category was significantly higher in the flat task (*t =* 3.8, *p* = 0.001). In other words, in each task, left lPFC privileged the abstract coding of a small number of task variable.

In the pAC (Fig 4C, right panel), as expected, CCGP was above chance for the auditory feature (flat task: *t =* 3.2, *p* < 0.001; hierarchy task: *p* < 0.001), which is consistent with pure selective coding. No other feature was abstractly coded in either task (*p*s > 0.1).

Collective, analysis of the cross-generalizability of lPFC representations showed that abstract coding is observed in a very small set of dichotomies, one of which reflects an important task variable. Crucially, in support of the task-tailored geometry hypothesis, different task-relevant dimensions were coded in abstract format across the two task structures. Having established these characteristics of lPFC control representations that differentiate it from the control region, pAC, we now direct our focus on lPFC and characterizing the geometry of the control representation across the two tasks more specifically.

### Local structure of lPFC representational geometry of the flat task shows high separability with no evidence for abstraction

The dominant global feature of the lPFC representation of the flat task is the coding axis encoding the XOR response categories. Patterns associated with each response category tend to cluster together on respective sides of a hyperplane separating the categories along this coding axis. We further characterized the local structure of these clusters, asking whether it is low or high-dimensional and if it is aligned across the clusters. To this end, we estimated hemodynamic response patterns in left lPFC parcels for 16 trial types, defined by the three stimulus features and the motor response mapping (i.e., relative location of the symbols). These were separated into two sets based on which response category they were associated with. For each set, we trained a set of linear classifiers to decode all identifiable task variables, separability and abstraction. And, finally, we asked whether these representations were aligned across the two clusters.

Across the two category clusters, 14 of 70 possible dichotomies were decoded above chance levels after multiple comparison correction for the number of dichotomies. Decoding accuracies (Fig 5A) were significantly above chance levels for all identifiable task-relevant dichotomies for both, the response category 1 cluster (visual feature 1: 54.5%, *p* < 0.001*; visual feature 2: 54.5%, *p <* 0.001*; auditory feature: 53.4%, *p* < 0.001*) and the response category 2 cluster (visual feature 1: 53.5%, *p* < 0.001*; visual feature 2: 52.9%, *p =* 0.003*; auditory feature: 53.5%, *p* < 0.001*), barring the motor response which was not decodable for either response category (all *ps >* 0.1). There were no significant differences across the response category clusters (*p*s > 0.1). Moreover, none of these local representations were abstract (Fig. 5C), including those encoding task variables with CCGP not reliably above chance for any of the dichotomies (all *p*s > 0.1). These results show that the representation of the flat task in the left lPFC is organized locally for high separability at the cost of generalizability.

**Figure 5.**
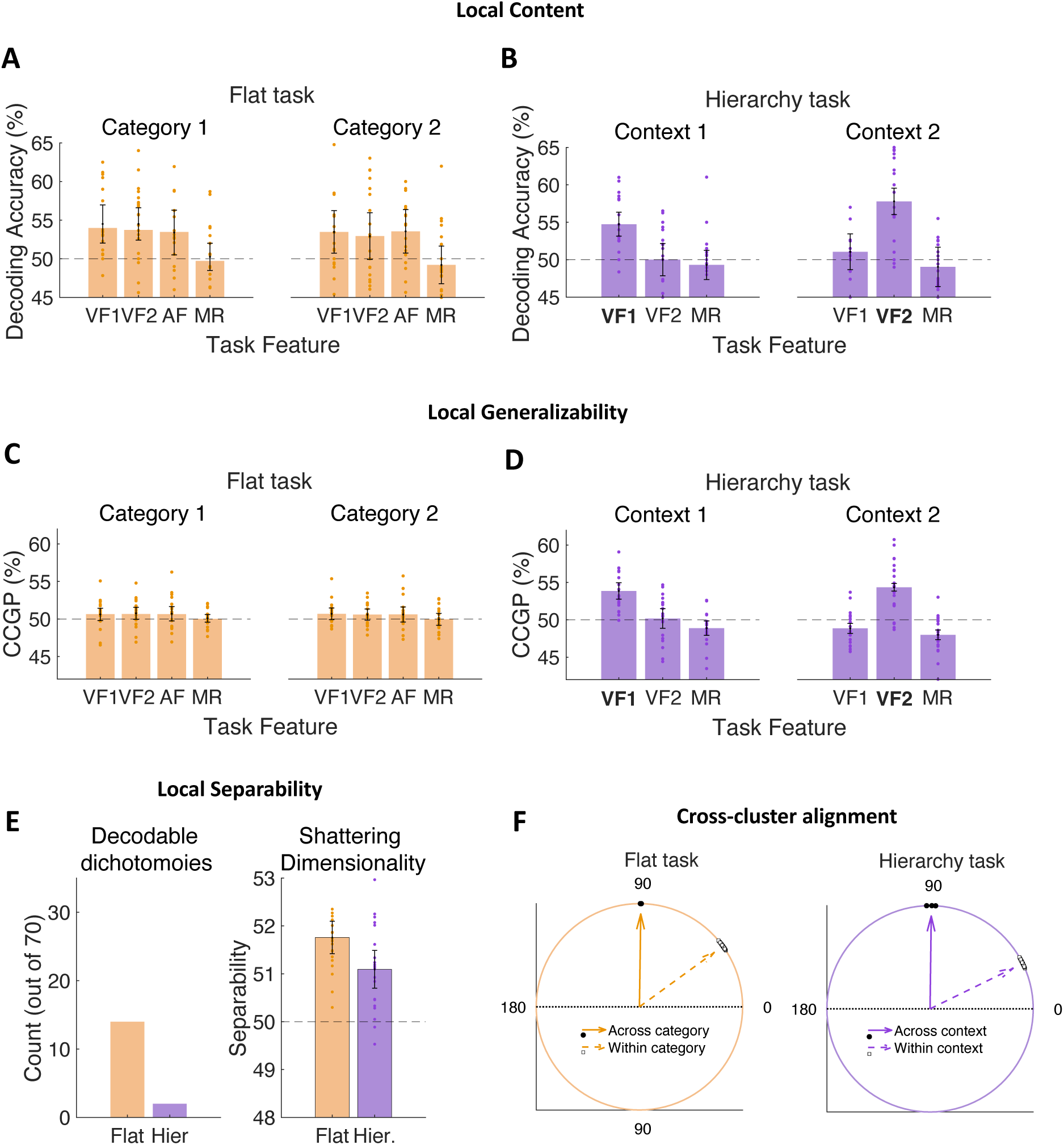
Properties of the local representation in the flat (orange) and hierarchy(purple) task. Local representational content was strikingly different across the task structures **(A)** In the flat task, all task variables could be decoded above chance levels in each category cluster. **(B)** In the hierarchy task, only the context-relevant stimulus feature is decoded above chance levels in each context cluster. **(C)** In the flat task, CCGP was at chance levels for all task variables. **(D)** In the hierarchy task, CCGP was above chance for the context-relevant stimulus feature in each context. **(E)** Local separability was starkly different across the two task structures. Assessed at the group-level with permutation tests, 14 dichotomies were linearly separable in the flat task while only 2 were linearly separable in the hierarchy task (E, left panel), with a paired t-test confirming greater separability in the flat task (*t_19_* = 2.8, *p* = 0.006). Local shattering dimensionality (E, right panel) was also significantly higher in the flat task (*t_19_* = 2.67, *p* = 0.015). **(F)** Local representations were not aligned across clusters in either the flat task (F, left panel) or the hierarchy task (F, right panel). Mean angles between coding axes for the same variable across clusters were close to orthogonal and significantly higher than the within-cluster angles. Mean angles pool across all decodable variables. All error bars reflect 95% confidence intervals.

Finally, we tested the extent to which the axes coding different task variables were aligned across the two context clusters. If each context is encoded in distinct subspaces, then these axes are predicted to be orthogonal. We trained the classifier to decode each dichotomy on the patterns of one response category and tested it on the patterns from the other category. Above-chance performance on the cross-generalization would demonstrate aligned coding, indicating some degree of representational overlap, while a failure to cross-generalize would be consistent with orthogonal coding. We also directly estimated the angle between the coding axes for each task variable across the two contexts.

Cross-category decoding accuracies were significantly above chance levels only for visual feature 2 (51%, *t =* 2.4, *p* = 0.025, BF_10_ = 2.4). The estimated mean angle between the respective coding axes across the two category clusters was 89.7 degrees (Fig. 5F, left panel) and not significantly different from orthogonal (p > 0.1) with the Bayes factor indicating moderate evidence in favor of the null (BF_10_ = 0.31). Therefore, there appeared to be no evidence for representational alignment across the two contexts.

In summary, these results build up a detailed picture of the representational geometry of the flat task in the left lPFC. Globally, this geometry consists of two clusters separated along a dominant axis that encodes the XOR response categories. Within this global picture, each cluster is locally organized into a geometry which supports linear readout along many dimensions including those that do not encode the main task variables, but not in abstract form. The clusters representing the two response categories appear to be independently organized such that axes encoding the same dimension are not always aligned across the two clusters.

### Local structure of left lPFC representational geometry in the hierarchy task is low-dimensional and recapitulates task structure

The dominant global feature of the lPFC representation of the hierarchy task is the coding axis encoding the latent context signaled by the auditory dimension. Patterns associated with each latent context tend to cluster together on respective sides of a hyperplane along this coding axis. We further characterized the local structure of these context clusters, as in the case of the flat task.

Across the two contexts, only 2 of 70 possible dichotomies were decodable above chance levels after multiple comparison correction for number of dichotomies. For both context clusters, decoding accuracies (Fig. 5B) were reliably above chance only for the context-relevant stimulus feature (context 1: 54.7%, *p* < 0.001; context 2: 57.8%, p < 0.001), and not for the context-irrelevant stimulus feature (context 1: 51.0%, *p* = 0.12; context 2: 50.0%, p = 0.49). A paired *t*-test confirmed stronger coding of the context-relevant feature compared to the context-irrelevant feature (*t =* 5.1, *p* < 0.001). Moreover, the local representation of the context-relevant stimulus feature was abstract, with CCGP (Fig. 5D) being reliably above chance (51.3%, *t* = 5.8, *p* < 0.001). Note that the context-relevant stimulus category is confounded with the response category and this finding could also be interpreted as the coding of the decision.

Finally, we tested whether the axes encoding the context-relevant stimulus feature across the two contexts were aligned or orthogonal. Cross-context decoding accuracies were not significantly above chance levels (49.8%, *p* > 0.1). The estimated angle between the respective coding axes across the two contexts was 89.8 degrees (Fig. 5F, right panel) and not significantly different from orthogonal (p > 0.1) with the Bayes factor indicating moderate evidence in favor of the null (BF_10_ = 0.25). Therefore, there appeared to be no evidence for representational alignment across contexts, suggesting that the lPFC learns orthogonal representations of the context-relevant stimulus features across the two contexts.

In summary, the hierarchy task was represented in lPFC with a geometry that recapitulated its hierarchical structure. Globally, this geometry consists of a dominant axis that encodes the higher-level context in an abstract, generalizable format, thus producing distinct contextual clusters. Within each cluster, the local geometry preferentially encoded only the context-relevant stimulus dimension and did so in an abstract, generalizable format. These subspaces appear to be independently organized such that axes encoding the same stimulus dimension may not be aligned across subspaces.

### Confirmatory analysis with Representational Similarity Analyses

We confirmed the detailed picture of the task-tailored geometries of each task built up over individual decoding analyses with representational similarity analysis (RSA) that permits a more global assessment ^52^. For both left and right lPFC, and separately for each task, we estimated a representational dissimilarity matrix (RDM) of the pair-wise, cross-validated Mahalanobis (‘crossnobis’) distances^53^ between the multi-voxel patterns associated with each of the eight trial types. Using multiple linear regression, we estimated the contribution of model RDMs that predicted pairwise dissimilarities based on different features of the representational geometry. This approach ensured that we tested the unique effects of each task variable, which is difficult to do with decoding analyses.

For the flat task, we tested model RDMs (Fig. SF1) that predicted pair-wise distances based on the three stimulus dimensions and the response category (Fig. SF2A). Confirming the results from the decoding analyses, in the left lPFC we found evidence for coding of the response category (*t =* 3.1, *p =* 0.006*) which explained unique variance in the pattern distances over and above the effects of the stimulus feature RDMs. We also found evidence for the coding of visual feature 2 (*t =* 2.6, *p =* 0.016*).

For the hierarchy task, we tested model RDMs that predicted pair-wise distances based on the three stimulus dimensions, the response category, or a context-dependent representation that preferentially encoded the context-relevant stimulus feature (Fig. SF2B). Confirming the results from the decoding analyses, we found evidence that both the auditory dimension/context (*t =* 10.9, *p <* 0.001*), the response category (*t* = 3.1, *p* = 0.005*), and the context-dependent representation of the relevant feature (*t =* 4.0, *p* < 0.001*). Therefore, the RSA confirms both the global and local features of the task-tailored representational geometry in the hierarchy task.

Finally, we also employed multi-dimensional scaling to employ the representational geometries consistent with group-level RDMs in each task (Fig. SF3). This analysis also confirmed the main features of our findings.

### Neural pattern variability along task-relevant coding axes correlates with behavior

We next asked whether the task-tailored geometries we identify in left lPFC shape task performance. To this end, we employed mixed-effects regression to test whether trial-by-trial variability in behavior (choice and RT) was correlated with neural pattern variability along task-relevant coding axes identified in decoding analyses. We estimated single-trial neural patterns in left lPFC for both the flat and hierarchy task and trained single-trial classifiers to decode task features in both tasks. We projected single trial patterns from left-out data onto the coding axes of these trained classifiers to obtain single-trial signed estimates of distance from the classification boundary. These signed distance estimates reflect the component of neural variability along the coding axis for the particular variable. If these neural components influence behavior, greater positive distance should be correlated with faster response times or more accurate responses. In the flat task, we focused on neural components coding the three stimulus features and the response category. In the hierarchy task, we focused on neural components coding the context feature, the context-relevant stimulus feature and the context-irrelevant stimulus feature.

In the flat task, the signed distance from the response category classification bound was the only regressor that explained variance in both response times (*β* = −0.004, *p* = 0.001, Fig. 6A) and choice (*β* = 0.09, *p* < 0.001, Fig. 6B). Therefore, patterns further from the classification bound along the response category coding axis were correlated with faster and more accurate performance.

**Figure 6.**
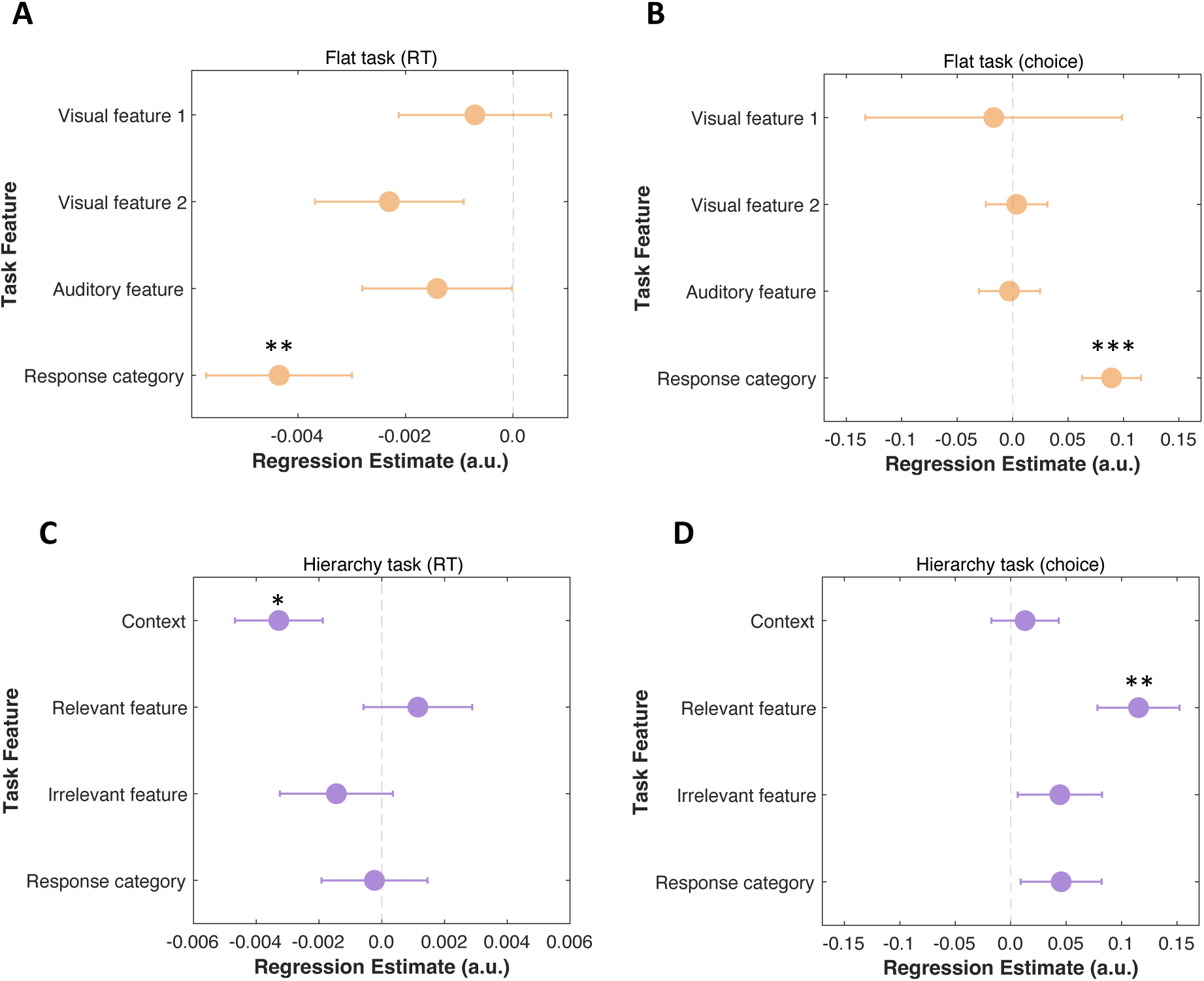
Task-relevant components of neural pattern variability in lPFC correlates with behavior. Results of mixed-effects regression of trial-by-trial response times and choices on trial-by-trial estimates of signed distances from hyperplane obtained for each task-relevant feature for the flat (A, B) and hierarchy (C, D) tasks are presented as forest plots. In the flat task, trial-by-trial variability in both response times (A) and choices (B) was explained uniquely by neural variability in the coding of response categories in left lPFC. In the hierarchy task, trial-by-trial response time variability was explained by variability in the coding of context information (C) while choices were explained by variability in the coding of the context-relevant stimulus feature (D). Error bars reflect standard error. * < 0.05; ** < 0.01; *** < 0.001.

In the hierarchy task, the signed distance from the context classification bound explained variance in response times (*β* = −0.003, *p* = 0.019, Fig. 6C), and the signed distance along the local, context-specific coding axes for the context-relevant stimulus features explained variance in choices (*β* = 0.32, *p* = 0.03, Fig. 6D). Therefore, improved performance was associated with stronger coding of context and the context-relevant stimulus feature.

In summary, these brain-behavior correlations support the behavioral relevance of these geometric features of lPFC representations, on a trial-by-trial basis. Further, they provide evidence that neural activity on the task-tailored manifolds for each task are a common determinant of behavior.

## Discussion

Collectively our results offer strong evidence that the lPFC learns task-tailored representational geometries to accommodate different tasks. In this case, lPFC representations were shaped differently to follow flat vs hierarchical task rule structures. Further, comparison of these different geometries provides evidence for the general principles that shape control representations in the lPFC. We elaborate these observations below.

In the flat task, combinations of three inputs were mapped, in a non-linear/XOR fashion to separate response categories. Correspondingly, the lPFC representational geometry was defined by a global axis abstractly encoding the XOR response categories (i.e., the outputs needed to select a motor response). Within the subspace defined by each response category, we observed a local geometry in which several task variables and their mixtures were linearly separable, with no evidence of local abstraction. Moreover, these local representations of task variables were not aligned across the subspaces. In other words, the lPFC representation geometry in the flat task was characterized by partial compression that afforded abstract coding of the task-relevant response categories. Sensitivity was maintained to changes in individual stimulus features, but only within the context of each response category.

This task-tailored geometry strikes a compromise between generalizability and separability. The global axis of the geometry accords with the demands of the flat task, where the primary basis for any two inputs being similar or not is their respective arbitrary membership in one or the other categories. Mapping inputs onto a one-dimensional manifold that only encodes the response categories provides for a robust readout of the required task output.

Indeed, the local structure observed within each response cluster is not strictly required for the downstream readout of the response category; a fully compressed global axis with no separation among individual stimulus inputs could support responding. So, why is this higher-dimensional local structure evident? Of course, one possibility is that this information is used for controlling other aspects of the task than responding, such as top-down attentional signals to perceptual processing and categorization. As such, this local structure may be shaped as much as the global axis. Though why these putative stimulus-level control signals would differ as a function of the response category is not immediately clear, which limits this interpretation.

Another possible account of this local structure is that it reflects the vestiges of an expressive, task-agnostic representation. Participants may initially represent the task inputs on a high-dimensional manifold during the initial stages of learning the task and then learn to compress to a one-dimensional manifold with experience. Though, as elaborated below, the system is biased to preserve expressivity due to how it discovers structure.

In the hierarchy task, the lPFC representational geometry fully recapitulates the structure of the hierarchy task. In this case, the geometry was dominated by a global axis abstractly encoding the auditory stimulus dimension, which signaled the superordinate rule context (e.g., attend to face versus attend to scene). Though we did not use different input domains than auditory to cue the context, we interpret this axis as likely representing the context rather than the auditory modality per se. Not only is this interpretation in line with previous findings^41^, it is also the most parsimonious account of our own data. In the flat task – wherein the auditory dimension did not act as context – there was no such organization around this auditory feature. It was only when auditory inputs cued the context, during the hierarchy task, that this axis of organization was evident.

This context coding axis defined context-dependent subspaces within each of which a locally low-dimensional geometry abstractly encoded only the context-relevant stimulus feature necessary for selecting a motor response. There was no evidence for the coding of the context-irrelevant stimulus feature within the contextual subspaces. Moreover, we found no evidence that the representation of the context-relevant stimulus feature was aligned across the two contexts, consistent with the representations in the two contexts being orthogonal^32,54^. This suggests that the lPFC employs separate context-specific readouts to identify the critical task-relevant output required to select a motor response, while minimizing the influence of potentially conflicting context-irrelevant inputs.

lPFC representational geometry we observe in the hierarchy task is consistent with prior studies in both macaques and humans, though we note that no prior study has compared the representational geometry of multiple tasks in the same participants. Roy et al. (2010) observed preferential coding of context-relevant stimulus categories, with largely segregated populations encoding the categories across the two contexts^25^. Flesch et al. similarly observed orthogonal, low-dimensional representations of the context-relevant feature with compression of the context-irrelevant feature in a broad frontoparietal cortex network, also demonstrating that such a geometry emerges in neural networks trained on a hierarchically structured task in a rich learning regime^32^. More generally, orthogonal coding has been consistently observed in lPFC in task settings with potential for interference, such as between the coding of targets and distractors^54^, and between previous and current task states^38^, suggesting a specific role for lPFC in encoding orthogonal subspaces in the service of cognitive control^36^. This accords with the proposal of “flexibility-via-subspaces” that has emerged in the literature on motor control^55^.

While our observation of local compression of irrelevant inputs in the hierarchy task is consistent with prior observations across species, our results do diverge in some respects from at least one observation in the non-human primate. Specifically, Mante et al. reported robust coding of both context-relevant and context-irrelevant stimulus information in the frontal eye fields (FEF) of macaques, with no evidence for compression^27^. In their study, context-dependent selection occurred via a mechanism operating through dynamics that enabled input from the relevant feature to push neural activity along a common response axis across contexts. While such a common response axis has been also observed in the lPFC^31^, we found no evidence for aligned coding of the response category across contexts in the hierarchy task, in line with other human studies^32^. This discrepancy may reflect the fact that training primates on such tasks typically involve a compositional shaping process where different components of the task are progressively introduced, with the response-only component introduced early before the categorizations are introduced. This may encourage the learning of a shared response axis. On the other hand, our participants were given verbal instructions for all task components in one go, with the differences across the two contexts emphasized.

lPFC representations across the two tasks showed no significant differences in the number of decodable dichotomies or shattering dimensionality at the global level. At first glance, this may seem surprising given the two very different task structures and the opportunity for abstraction in the hierarchy task where only one of the two visual stimulus features were relevant on a given trial. However, this is to be expected. First, it should be the case that *global* shattering dimensionality does not differ across the two task structures. By design, both tasks require a non-linear computation with the inputs to arrive at the response. Therefore, both tasks show a similar (intermediate) dimensional representation which helps produce separate subspaces in each task representation. Second, at the local level, within each subspace defined by either context (hierarchy task) or response categories (flat task), we do find a striking difference in dimensionality across the two task structures. In the hierarchy task, we indeed observe a lower dimensionality within each context-specific subspace, with local geometry mainly reflecting the coding of the context-relevant stimulus feature. On the other hand, in the flat task, we observe an expansion of dimensionality within each response category subspace, with local geometry showing higher separability. These observations are consistent with tailoring of the geometry, including its dimensionality, as a function of task structure.

Crucially, the task-tailored geometries we observed in the lPFC were linked to participants’ behavior in the task. Across both the flat and hierarchy task, a portion of the trial-by-trial variance in response times and choices was explained by trial-by-trial variance in coding of key task variables in lPFC. Importantly, rather than general variability in neural activity, components of variability along specific task-tailored coding axes explained behavior. In the flat task, variability in the coding of response categories, but not the individual stimulus features, explained behavioral variability. In the hierarchy task, neural variability along the coding axis for context and the context-relevant (but not context-irrelevant) stimulus feature explained behavioral variability. These specific brain-behavior relationships provide evidence that the geometric features of task-tailored lPFC representations, such as their organization around different coding axes, are consequential for successful, controlled behavior.

While the task-specific geometries we have uncovered here are consistent with previous observations of task-sensitivity of lPFC neural activity, and a few studies examining lPFC representational geometries in individual tasks, our study is unique in that we tested the representation of these different task structures in the same participants, allowing us to directly address the question of how differently structured tasks are accommodated in the human lPFC. In particular, our study goes beyond prior work documenting task sensitivity by demonstrating that in each task, lPFC neural activity is organized into task-tailored geometries that facilitate or enhance the specific computations or readouts needed for the particular task. In the flat task, our decoding and CCGP results show that the shape of the task manifold enables efficient and robust readout of response categories, which is the key variable for selecting responses. In the hierarchy task, we observed both separation into contextual subspaces, and distinctly shaped local manifolds that separated along the context-relevant feature, but collapsed along the context-irrelevant feature enabling efficient and robust readout of the response category in each context. Therefore, lPFC representational geometry in each task was shaped to enhance task-specific computational demands. This is further corroborated by the observed relationship between behavioral variability and neural variability along the same components of the task-tailored geometries.

Our empirical resolution of the question of whether the lPFC employs task-tailored or task-agnostic representations has implications for the broader understanding of the regions’ function in flexible behavior. One broad idea is that the function of lPFC is to provide an ‘adaptive’ representational resource. On this view, lPFC neurons can rapidly assemble into ensembles to help us perform tasks that we do not have much experience with^42^. To perform such a function, a task agnostic representational format may be suitable^34,35^. Another broad idea is that the lPFC, like cortical regions throughout the brain, are shaped by the statistical structure of experience, learning and storing a variety of representations, and extracting their compositional structure^3^. Such a view would predict that experience with a task would shape lPFC representations^56^. Our results clearly show that, at least after substantial experience with a task, lPFC representations are tailored to the structure of the task. Therefore, our results support the latter, statistical learning view of lPFC function though it remains possible that during the early phases of performing a task, lPFC representations are more task agnostic^40^.

While we found clear evidence for distinct, task-tailored representations for the two very different task structures we studied, even more notable were the striking similarities in the general representational strategy employed by lPFC across the two tasks. The tasks did not differ either in what task variables were decodable, the overall separability, mixed-selective separability, or the overall generalizability of the lPFC representation. For both task structures, more than half of the possible dichotomies were linearly separable. And, across both task structures, separability was preferentially enhanced for dimensions encoding task variables at the cost of other dimensions. Therefore, beyond the characterization of the detailed structure of the task-tailored geometries, our results comparing the two task structures illuminate the general representational strategy lPFC employs, highlighting a set of key principles governing the lPFC neural code.

First, lPFC preferentially encodes a diversity of environmental variables that are relevant to the task at hand, while suppressing orthogonal, task-irrelevant variables. This deviates strongly from the coding principle of putative input regions, like the primary auditory cortex, which obligatorily encodes auditory variables in the environment, regardless of their task-relevance and shows no evidence of mixing with other input features. This principle of diverse lPFC coding accords with the well-replicated finding that lPFC neurons code for information about whatever task is currently being performed.

Second, lPFC representations are not obligatorily low or high dimensional, but can reflect a range of possible compromises between a maximally compressed, pure-selective code and a maximally expanded, highly conjunctive code based on the needs of the task^35^. This flexible representational format enables learning processes to produce a variety of task-tailored geometries. Indeed, for most tasks, lPFC representations will be of intermediate dimensionality because it affords linear separability for several task variables and their mixtures, along with abstract, generalizable coding of a small number of critical task variables.

What variables are selected for abstract coding? In our study, across both task structures, the key output variable was coded abstractly. We suggest that one key driver for abstract coding is the need for robust, generalizable readout. In the flat task, the response category was coded abstractly. An abstract representation of the task output allows for the same geometry to be re-used in novel situations where the response categories are preserved, as has been observed in macaques^40^ and humans^57^. In the hierarchy task, the context-relevant stimulus dimension was coded abstractly at the local level. Not only does this again fully determine the response category, compression over the irrelevant dimension makes it more robust to a salient source of distraction.

Third, the lPFC organizes neural responses into orthogonal subspaces. Across both task structures, we observed distinct clusters of neural responses – in the flat task these were organized by response category, while in the hierarchy task, these were organized by context. Each cluster showed a unique local organization which did not align with that of the other cluster, strongly suggesting orthogonal coding.

An advantage of such an organizational scheme is that it affords protection for coded information from interference. In the hierarchy task, a key source of conflict is that between the context-relevant and context-irrelevant stimulus features which can drive different responses. By organizing neural responses of each context into orthogonal subspaces, the lPFC could selectively code only the context-relevant stimulus feature in the relevant contextual subspace, suppressing the context-irrelevant information and thus minimizing interference. Indeed, orthogonal coding has been widely observed in macaques^15,16,25,58–60^ and human lPFC^32,38^ and has been implicated in a variety of settings that demand the minimization of interference.

Fourth, and more speculatively, the lPFC may combine a default preference for high-dimensional coding in the lPFC with learning-driven dimensionality reduction for discovering structure. A surprising novel finding in our study is the strikingly different local organization we observe across the task structures. As discussed above, the local structure in the hierarchy task recapitulates the lower-level structure of the hierarchy task where only one stimulus feature is context-relevant. On the other hand, it is less clear how orthogonal coding of locally high-dimensional structures is related to the demands of the flat task. Indeed, the flat task has no obvious sources of interference that would be reduced with this subspace organization and no obvious requirement for high separability at the local level.

One possible account of these findings is that orthogonal coding may be a general property of lPFC coding, arising not from the tailoring of the representation to the task, but from a default preference for high-dimensional coding. On this view, the lPFC would initially always encode its inputs on a high-dimensional manifold, where every response is orthogonally coded, with learning processes reshaping the manifold, retaining separability only along dimensions that the task requires. The hierarchy task has both global (i.e., mappings are consistent within each context, but different across each context) and local (i.e., only one feature is relevant within each context) structure that can be discovered by a learning process, while the flat task had only global structure and no local structure. Therefore, the lPFC responses in the hierarchy task were organized in a locally low-dimensional organization which privileged the coding of the context-relevant feature, while the flat task remained locally high-dimensional.

Several studies in macaques are consistent with this possibility. Wojcik et al. in their recent study found that lPFC representations are high-dimensional at the onset of learning, being slowly shaped to be task-tailored with experience^40^. Bernardi et al. found evidence for representational geometries that incorporated abstract coding by minimally altering a high-dimensional code. Finally, high-dimensional coding has been observed in the lPFC when the task itself does not impose a low-dimensional structure^34^.

An account that combines a natural preference for high-dimensional coding in the lPFC with learning-driven dimensionality reduction for discovering structure in tasks may explain how the lPFC resolves the separability vs generalizability tradeoff. Encoding novel inputs on a high-dimensional, task-agnostic manifold by default allows sensitivity to a variety of potentially relevant task variables in the initial stages of learning, supporting expressivity. With learning, this manifold may be reshaped to make it task-tailored by enhancing both the separability and generalizability of key task variables while reducing them for others. Such task-tailored representations would then support generalization to other tasks with similar structures as has been observed in macaques^40^ and humans^57^.

Such generalization would dramatically speed up learning in novel settings and may explain why, in our study, participants learned the hierarchy task much more rapidly than the flat task. For the hierarchy task, participants were likely able to re-use existing representations of rules learned in other contexts. On the other hand, given the contrived nature of the XOR classification rule in the flat task, they may have had to build the representation from scratch.

This is in line with accounts of faster hierarchical rule learning in a more traditional reinforcement learning setting, as well^10,61^. Given that we did not directly measure the neural representations in lPFC during the early stages of learning, however, we cannot directly test these predictions. Doing so will be an important direction for future research.

Our study has some limitations. The use of fMRI constrains our ability to examine within-trial dynamics in the representational geometry. Prior work using electrophysiology in macaques^16,26,62,63^, and using EEG in humans^64,65^, has documented marked shifts in the representational geometry within a trial on the order of hundreds of milliseconds. Our estimates of the representational geometry, therefore, reflect an aggregate measure of this complex trajectory. Several recent EEG studies have documented the importance of conjunctive representation of context, stimulus and response as being critical for the timing, accuracy and control of context-dependent action selection, though their locus in the brain is presently unknown^64–67^. Our results in the hierarchy task accord with these observations in that we observe context-dependent coding of the relevant stimulus (i.e., a context-stimulus conjunctive code). A recent EEG study documented a transient expansion of representational dimensionality in the moments before successful response selection which coincided with stable coding of conjunctive representations^65^. Given the relatively high separability and the context-dependent nature of the coding we observed in the hierarchy task, one possibility is that our fMRI models capture this moment in the representational dynamics though, we cannot be sure of this possibility with only BOLD fMRI measurements.

Another limitation related to fMRI concerns the notoriously noisy lPFC BOLD measurements^50^. To maximize power for detecting multivariate effects in lPFC, we employed a deep sampling strategy^68–72^ and restricted our sample to participants who could achieve a high level of accuracy and speed in both the flat and hierarchy task within a fixed training period. Therefore, our results only apply to other similar, highly-trained and high-performing participants and may not generalize to participants who take longer to learn the task or use a less effective strategy to respond, particularly in the more difficult flat task. This leaves open important questions about individual differences. For example, would participants who take longer to learn to represent the task with a qualitatively different, poorly adapted geometry, or would they employ an inefficient version of the same geometry as what we identified? Nonetheless, our results reveal that the lPFC can accommodate tasks of different structure with task-tailored representational geometries.

Finally, our study was focused specifically on the coding principles of lPFC, which we defined as parcels from a single functional network^47^. We have not examined other lPFC parcels that have been localized to distinct functional networks. This leaves open the question of whether representations in other networks within lPFC may be driven by different organizing principles, and how they might collectively interact to support controlled behavior. More broadly, our study raises the question of whether these principles generalize beyond lPFC, and if so, how broadly. The whole-brain fMRI data we have collected provide ample opportunities to explore the principles of coding across the brain.

In conclusion, we tested two disparate accounts of how the lPFC accommodates tasks of different structures, finding clear evidence for task-tailored representations of task-relevant environmental variables in well-trained human participants. By comparing and contrasting the representational geometries in lPFC across two disparate task structures, we identify four key principles that govern lPFC control representations: i) diverse coding of task-relevant inputs, ii) a flexible representational format that supports the coding of task-tailored representations, iii) orthogonal coding that reduces interference, and iv) a default preference for high-dimensional coding in the lPFC with learning-driven dimensionality reduction for discovering structure.

## Methods

### Participants

All participants in the main experiment were right-handed; had normal or corrected-to-normal vision and hearing; and were screened for the presence of psychiatric or neurological conditions, psychoactive medication use. Participants who participated in the fMRI scanning phase were additionally screened for contraindications for MRI. All participants gave their written informed consent to participate in this study, as approved by the Human Research Protections Office at Brown University. Participants were monetarily compensated for their participation at each session and received a bonus payment if they completed all 11 scanning sessions.

A total of 94 participants (58 female, 33 male, 3 declined to answer; mean age = 22.9 ± 4.7) were recruited during the behavioral training phase of the study. In this phase, participants learned the two tasks (detailed below) and performed a series of practice blocks, with the instruction to maximize performance. Those who achieved the *a priori* performance criteria of > 85% overall performance with >80% performance on each of the 8 trial types within 4 days of training for each task were invited to participate in the scanning (fMRI) phase of the study.

Out of the 94 total participants, 21 withdrew from the study before completing the training phase, and an additional 37 did not meet performance criteria on at least one of the tasks. The remaining 36 participants were invited to the subsequent scanner phase. Of these, 3 participants were excluded due to contraindications for MRI, and 13 participants either withdrew before completing all sessions or were withdrawn for failing to comply with instructions. 20 completed the full study and are included in the analyses presented here (14 female, 6 male, mean age 22.4 years, SD = 4.5 years).

### Stimuli

Participants performed two categorization tasks that mapped multidimensional stimuli consisting of 3 binary features onto abstract categories symbolized by simple shapes. The three stimulus features included two naturalistic images and an auditory clip of a spoken number or word. Two stimulus sets were employed for each participant, one for each of the tasks. One set consisted of naturalistic images of faces (adult or child) and scenes (indoor or outdoor), along with auditory clips of spoken numbers (low or high). The other set consisted of naturalistic images of animals (birds or mammals) and objects (edible or inedible), along with auditory clips of spoken words (nouns or verbs). The mapping of stimulus sets to tasks (hierarchy or flat as described below) was counterbalanced across participants. The same set was always used for a given task within-participant.

A total of 13,600 naturalistic images (faces, scenes, animals, objects) and 96 audio clips (spoken words and numbers) were employed across the study. During the behavioral training phase, trial-unique images were employed within a single block, but images could be repeated across blocks. In the scanning phase, the images were trial-unique across all blocks such that no image was presented twice during the entire scanning phase across all days. However, images in the scanning phase were sampled from the underlying set, and therefore may have been seen by participants during the training phase. On average, each image was used on 1.6 trials across the entire study per participant. Given the noisy environment in the scanner and the need to use high-quality auditory stimuli, audio clips were not trial-unique but included multiple speakers producing each word/number. Each audio clip was used on average twice in each scanning run and the same set of auditory clips were repeated across runs. On average, each clip was used 91 times across the entire study.

#### Stimuli

Stimuli consisted of two visual images and an auditory, spoken word. All three stimulus components were systematically manipulated along two orthogonal dimensions, one of which was relevant to the tasks and one was irrelevant. Two stimulus sets were used across the study which are detailed below.

#### Stimulus set 1

*Face Images:* Face images were systematically selected to vary along two orthogonal dimensions of age (child vs adult) and hair length (short vs long). Only the child vs adult dimension was relevant for either task. Simulated face images were first generated using StyleGAN2 (Karras et al., 2019). These faces were then mapped to the binary categories of adult (defined as someone judged to be over 35) or child (defined as someone judged to be under 18). Orthogonally, they were mapped to categories of people with either short hair or long hair. Age mappings were validated by independent raters (N=15, mean age 31.6, SD 2.1, 9 male, 6 female) on Amazon Mechanical Turk who made forced-choice child vs adult decisions about each image. Images were only included in the final stimulus set if at least two researchers and two MTurk participants agreed on the age category. Participants were informed at the end of the experiment that the faces did not depict real humans.

##### Scene Images

Scene images were systematically selected to vary along orthogonal dimensions of indoor vs outdoor and scenes with vs without people. Only the indoor vs outdoor dimension was relevant for the tasks. All scene images were obtained from the Places205 dataset which contains naturalistic scenes from 205 different categories^73^.

##### Number Audio clips

Spoken number stimuli were systematically created to vary along orthogonal dimensions of magnitude and parity. Thus, numbers could be low vs high, as well as odd vs even. Only the low vs high dimension was relevant for either task.1s long audio clips contained spoken number words “three”, “four”, “eight”, and “nine”. Clips were generated by an online text-to-speech converter (www.naturalreaders.com) using six different synthetic voices.

#### Stimulus set 2

##### Animal Images

Animal images were systematically selected to vary along orthogonal categorical dimensions of birds vs mammals and domestic vs wild, with only the birds vs mammals dimension being relevant to the tasks. Animal images were manually screened and assembled from a variety of online sources including the iNaturalist dataset (www.inaturalist.org), Pixabay (www.pixabay.com) and Google Images (images.google.com) to minimize ambiguity. Images were only selected if they unambiguously depicted one or two animals of the same type.

##### Object Images

Object images were systematically selected to vary along orthogonal dimensions of edible vs inedible and natural vs manmade. Only the edible vs inedible dimension was relevant to the tasks. Object images were manually screened and assembled from ImageNet^74^ and the THINGS database^75^ to minimize ambiguity.

##### Word Audio clips

Spoken word stimuli were systematically created to vary along orthogonal dimensions of noun vs verbs and words starting with the /d/ vs /l/ sounds. Only the noun vs verb distinction was 1s long audio clips consisted of the spoken words, “door”, “lake”, “lose” and “dig”. The stimuli were recorded in-house using a Logitech Blue Snowball microphone in a sound-proofed room.

In addition to trials with these complex stimuli, both tasks also featured infrequent “null” trials which presented two images of random black and white noise along with a pure 300 Hz tone. The noise images and the tone were not trial-unique.

### Task design

Participants learned to perform two categorization tasks with different structures, labelled as the ‘flat task’, and the ‘hierarchy task’. Each task required them to employ task-specific rules to select the category that one of 8 stimulus types (consisting of three binary features) belonged to, and then indicate their choice by pressing one of two keys. Across both tasks, trials followed the same structure (Fig. 1).

Stimulus presentation was implemented in PsychoPy^76^. Each trial began with the simultaneous presentation of two images side-by-side at the center of the screen along with the audio clip over headphones, for 1 s. This stimulus display provided the information needed to make the category decision based on the rules for that task. The left-right location of the two different image types (i.e., faces and scenes for stimulus set 1 or animals and objects for stimulus set 2) was randomized across trials and controlled such that each trial-type had an equal number of each possible configuration.

Simultaneously with the images and auditory clip, a response panel consisting of two shapes also appeared at the bottom of the screen to the left and right of the fixation cross (Fig. 1. The shapes differed by task. Specifically, a square and circle were presented for the flat task, whereas a triangle and pentagon were presented for the hierarchy task. Each shape represented one of the two possible categories and their position (left or right of fixation) on the screen indicated which key (left or right) the participant should press to indicate their selected category.

The left-right position of the two shapes varied randomly from trial to trial such that the position would be the same as the previous trial on approximately 50% of the trials, and each trial-type was associated with an equal number of left and right responses. This ensured that the participant’s motor response was not confounded with their categorization decision across trials. The response panel was presented for 2 s, which was also the window during which the participants had to respond. If participants pressed a key during the response window, the corresponding shape would turn gray.

Following the response period, there was an inter-trial interval (ITI) of 2-10s during which a white fixation cross was presented on a black background. During the behavioral training phase participants were also provided feedback. Specifically, the fixation cross would turn green or red for the first 500ms of the ITI to indicate correct or incorrect responses, respectively. During the scanning phase, participants did not receive feedback and the fixation cross remained white throughout the inter-trial interval.

While the same stimulus displays could be used for either task, the two tasks differed in their rule structures, leading to different responses on the same inputs across participants. Note that in both cases, participants were instructed about the rules explicitly, as described below. Nonetheless, they had to learn to use the rules efficiently, which required practice. We elaborate rules and procedures for each task below.

#### Flat Task

The mapping of stimuli to categories for the flat task is shown in Fig 1A (left panel). The mapping in the flat task was based on a latent, three-dimensional XOR or parity rule. Specifically, two stimuli which differ in an odd number of features belong to different categories, while two stimuli which differ in an even number of features belong to the same category. This ensures that for every trial, all three features are necessary to make a correct categorization and prevents any grouping within the rule structure, which can simplify the problem beyond two abstract categories. Importantly, however, this XOR rule was never described to the participants. Rather, they were only shown the mapping of each set of stimuli into the two categories and were asked to memorize these arbitrary mappings (see procedures below).

#### Hierarchy Task

The mapping of stimuli to categories for the hierarchy task is shown in Fig. 1A. The hierarchy task is organized around two contexts, defined by the auditory feature. The context determines which one of the other two stimulus features is relevant to the categorical decision. For example, a rule might be “if you hear a high number, the face determines the category: an adult face indicates the square category and a child’s face indicates the circle category.” So, while all three stimulus features were available on every trial and were relevant across the contexts, on any single trial only two are ever necessary to make a correct category decision. Note that one could, in principle, also perform the hierarchy task using the same memorization of stimulus groupings to categories used for the flat task. Nevertheless, the hierarchical rule was given explicitly to participants, and they were encouraged to follow a hierarchical strategy. This ensured that participants were oriented to different task structures.

Rules were presented at the start of each block during training and participants were encouraged to take as much time as they wanted to review the rules. They received veridical feedback on each trial during the training phase.

### Behavioral Training Protocol

During training, blocks contained 11 of each of the eight main trial types and the null events. ITIs were varied between 2 and 8 seconds. Training blocks took approximately eight minutes to complete.

Participants learned the two tasks sequentially outside the scanner. Behavioral training was carried out over up to four 90-minute lab visits during which the participant practiced as many blocks as time would allow. On average, participants completed 7 blocks per session, each lasting approximately 8 minutes. Participants returned for a minimum of 2 and up to 4 training sessions until they achieved a session accuracy of 85% overall and 80% accuracy on each of the eight individual trial types. On average, the 20 participants who completed all portions of the study required 2.7 sessions to learn the flat task and 2 sessions to learn the hierarchy task.

Participants were more likely to fail to reach the criterion on the flat task, which was harder to learn and perform. This necessitated a pragmatic choice of fixing training order with the flat task being always trained first to minimize participant dropout and associated cost. While it is possible that learning the flat task first influenced the subsequently learned hierarchy task, we believe this is unlikely given that the two tasks shared no stimuli within-participant and had very different task structures. Nevertheless, participants were always trained on both tasks before any scanning, and the order of the tasks was counterbalanced during the scanning phase. In the scanned sample, ten participants learned the flat task with the face, scene, and number stimuli and the hierarchy task with the animal, object, and word stimuli; while the other ten had the task to stimulus set mapping reversed.

### fMRI Task Protocol

Participants were scanned for a total of eleven days, an initial scan day for structural, resting state and localizer tasks, and 5 successive days each for the two tasks. We describe the typical protocol below and any deviations are detailed in the supplementary (Supplementary section S1). After the initial scan day, half of the participants (N=10; five using each stimulus set to task mapping) were scanned while performing the flat task first, and the other half were scanned while performing the hierarchy task first. All scans occurred between 7 and 11 am, and the ten task scans were performed within an average of 20 calendar days of each other (min=10, max=30).

Before entering the scanner, participants reviewed the rules and practiced two blocks of the task they would be performing in the scanner that day to refamiliarize themselves with the rules. One block included trial-wise feedback, and the other had no feedback. In each session, participants performed five blocks of the task while being scanned, yielding a total of 25 blocks per task which translates into a total of 1975 task trials and 200 null trials during the performance of each task. Participants could review the task rules before each block of the task but did not receive any feedback on their performance. After the study, participants were debriefed on their strategies and thoughts about the tasks using a structured interview.

Trial sequences during scanning were counterbalanced such that each of the eight possible trial types followed itself and every other type of trial at least once during a block (64 possible transitions), in addition to following null trials (8 additional transitions). To do this efficiently, we generated de Bruijn sequences using a published tool^77^. To create the full trial sequences, the final six transitions of each sequence were prepended to the start of their respective trial sequences. This process yielded sequences 87 trials long, with at least 9 instances of each of the eight main trial types in each block. A single block was presented during each scanning run.

Inter-trial intervals (ITIs) were samples from a truncated exponential distribution with a mean of two seconds and a maximum of 10 seconds, rounded to the nearest second. Given the trial transition and task length, we generated 100k possible timing sequences and then chose the 25 with maximum efficiency, calculated by averaging over all possible binary trial-type contrasts^78^. One trial sequence was used in each run and the mapping of trial sequence to run was randomized such that specific stimuli were not associated with specific trial sequences across participants. All scanning runs began with six seconds of fixation and ended with 16 seconds of fixation.

### fMRI Scanning Protocol

Whole-brain imaging was performed using a Siemens 3-T Magnetom PRISMA system with a 64-channel head coil. One high-resolution T1-weighted multi-echo magnetization prepared rapid gradient echo image (T1MEMPRAGE) was acquired as a structural image (repetition time (TR) = 2530 ms, echo times (TE) = 1.69, 3.55, 5.41, and 7.27 ms, flip angle = 7 degrees, 176 sagittal slices, 1 × 1 × 1 mm voxels). This image is used for all normalization procedures for the data in this paper. On at least four of the task days, a high-resolution T1-weighted magnetization prepared rapid gradient echo image was acquired; not further analyzed in this manuscript (TR = 1900 ms, TE = 3.02 ms, flip angle = 9 degrees, 160 sagittal slices, 1 × 1 × 1 mm voxels). Whole brain functional volumes were acquired using a gradient-echo sequence (TR = 1000 ms, TE = 32 ms, flip angle = 64 degrees, SMS = 5, 65 interleaved axial slices aligned with the AC-PC plane, 2.4 × 2.4 × 2.4 mm voxels). Each task functional run lasted 534s. Soft padding was used to restrict head motion throughout the experiment, and a vitamin D pill was placed on the right side of the participant’s forehead to verify left-right orientation. Stimuli were presented on a 32 in monitor at the back of the bore of the magnet, and participants viewed the screen through a mirror attached to the head coil. Participants used a five-button fiber optic response pad to register their button press responses (Current Designs, Philadelphia, PA).

Participants wore MR-compatible Avotech headphones. The sound volume was adapted for each participant at the start of each session. In addition to the BOLD signal, participants’ heart rates and respiration were measured during scanning sessions with an MR-compatible pulse oximeter and breathing belt (Siemens). In a total of 24 runs across 5 participants during the scanning sessions of the hierarchy task, and 25 runs across 6 participants in the flat task, technical difficulties (e.g., uncharged or broken equipment, improper participant placement/calibration) prevented the collection of physiological data. The status of this equipment did not interfere with the collection of MRI data.

### fMRI Preprocessing

Functional data were preprocessed using SPM12 (https://www.fil.ion.ucl.ac.uk/spm/). The quality of the functional data of each participant was first assessed through visual inspection and TSdiffAna (sourceforge.net/projects/spmtools/). Slice timing correction was carried out by resampling all slices within a volume to match the timing of a reference slice that was collected at 452.5 ms (i.e., closest to the halfway point of volume acquisition). Next, the effects of head motion during the functional runs were corrected with a three-step procedure. First, each volume in a run was registered to the first image in the run and individual run-mean was computed, and all run images were then registered to the run-means. using rigid-body transformation. Movement was assessed at this stage, within each run, to ensure that all volumes were within one voxel (2.4mm) of movement in all directions. No outlier volumes were detected under this criterion in the final sample. Next, these individual run-mean images were averaged to compute a global mean image across all runs. Finally, all volumes across all runs were registered to this global mean image. Anatomical images were also co-registered to this global mean. Data processed to this stage (slice time and motion corrected) were used for decoding and representational similarity analyses.

The anatomical T1MEMPRAGE image for each participant was normalized to MNI space and used to create a brain mask using the Brain Extraction Tool in FMRIB Software Library (fsl.fmrib.ox.ac.uk/fsl/fslwiki/).

### Regions of Interest

We derived our regions of interest (ROIs) from the 400 parcel map from the Schaefer parcellation^47^ (www.github.com/ThomasYeoLab/Standalone_Schaefer2018_LocalGlobal) which have been previously mapped to a set of 17 functional networks defined by resting state connectivity^48^, specifically using the ten lPFC ROIs mapped to the “Control A network” (parcels 128-132 and 330-334). Prior work on hierarchical tasks^5,79,80^ found that these regions are engaged in tasks similar to ours. Moreover, these regions were chosen because of their overlap with the multiple demand network which has also been identified in the macaque, where it includes the lPFC regions that provide the scientific premise for our hypotheses. Voxels across all lPFC parcels within each hemisphere were pooled to create single left and right lPFC ROIs which were employed for all analyses reported in the main text.

In addition, we derived primary auditory cortex ROIs from a term-based meta-analysis on Neurosynth (www.neurosynth,org) using an association test to obtain a map of brain regions preferentially related to the term “primary auditory” ^81^. This map was cleaned up to remove non-contiguous voxels outside the primary cluster and split into left and right auditory cortex.

All ROIs were transformed into each participant’s native space using SPM12’s reverse normalization procedure. Following native space transformation, ROIs were masked to include only voxels which have at least an 80% probability of being gray matter, based on SPM12’s unified segmentation for each participant. The average number of voxels in each ROI in each participant’s native space is provided in the supplement (Supplementary section S2).

### General Linear Modeling of fMRI data

Functional data were analyzed under the assumptions of the general linear model (GLM) using SPM12. Regressors were convolved with SPM’s canonical hemodynamic response function (HRF) with time and dispersion derivatives. Models were fit to voxel-wise timecourses for the whole brain level and were defined in the same way for both tasks. GLMs were separately fit to unsmoothed, native space data from each run.

We fit three separate GLMs. The goal of the first GLM (GLM1) was to estimate the activity related to each of the eight types of trials in each run separately for both tasks and to do so in as unbiased a manner as possible such that all eight estimates for a given run would be generated from equal amounts of trials and were balanced for confounding factors like associated motor responses. Specifically, all runs contained at least nine of each trial type, but if a participant made errors there would be fewer trials of certain types, leading to imbalanced amounts of data contributing to each activity estimate. That imbalance could bias decoding. Additionally, individual trial types could be associated with both left and right motor responses. An imbalance in the relative contribution of the two motor responses to different trial types could also bias decoding. To remove these biases, each of the eight trial types was randomly *sub-sampled* to the performance of the worst trial type and to ensure an equal number of left and right responses. For example, if a participant got eight trials of type A correct and five trials of type B correct, the model would sample randomly from five type A trials for that run. Additional correct trials (the three additional type A trials) would be modelled by separate trial-type specific ‘spillover’ regressors, as necessary. The same procedure was also used to balance motor responses associated with each trial type. These spillover regressors modeled the contribution of these trials to the voxel response, but they were not used for additional decoding or RSA analyses. The trials sampled to contribute to the main trial type regressors were chosen randomly, and the results of further analyses were insensitive to which random subset were chosen.

To summarize, then, for each run, GLM1 had eight main trial-type regressors, a single regressor each for all error trials and static trials, and as many additional spill-over regressors as necessary (between 0 and 7, with a mean of 6). Because of this conservative estimation approach, we dropped individual runs from additional analyses which would have fewer than four trials contributing to each of the main trial-type regressors. On average, 21.7 (min=11, std=4.07) runs contributed to regressions in the flat task and 23.8 (min=17, std=2.19) runs contributed to regressions in the hierarchical task. A mean of 6.6 trials contributed to each regressor in the flat task and 7.3 trials in the hierarchy task.

Two additional GLMs were built using the same procedure as above. One model (GLM2) estimated the responses associated with each of the 8-trial types but now split by what motor response mapping was employed, resulting in 16 trial-type regressors. Another model (GLM3) was identical to GLM1 except that it replaced the 8 trail-types regressors with 8 regressors for pseudo-trial-types defined by the orthogonal, task-irrelevant features of each stimulus. The subsampling procedure described for GLM1 was also reapplied in the construction of regressors for GLM2 and GLM3 to equate the number of trials and balance the motor responses contributing to each trial-type regressor in each run.

Finally, to estimate fMRI responses to single trials for analysis of brain-behavior relationships, we employed a Least-Square-Single (LSS) approach^82^. Briefly, for every subject and task, a suite of GLMs were built to estimate the response associated with each trial. Each GLM consisted of two regressors of interest, one for the trial whose response was to be estimated, and another regressor for all other trials. Therefore, each single-trial estimate came from a distinct GLM constructed for that particular trial. For simplicity, we refer to this suite of GLMs as GLM4.

All models otherwise included identical nuisance regressors. To reduce the impact of time-on-task on estimated regression weights, the duration of trial events in each of the above regressors was set to the trial-wise RT, or the duration of the stimulus display (2 seconds) for missed responses^83^. Nuisance regressors for left and right button presses with an onset at the time of participant response and a duration of 0 seconds were included. An additional nuisance regressor for the run mean was also included. Participant motion was captured in three translational and three rotational components, and these were used to generate a total of twenty-four motion-related nuisance regressors: absolute values, difference from prior volume, and square of each term^84^. All motion estimates were obtained within-run.

We did not employ high-pass filtering to reduce low-frequency noise. Instead, the first five principal components of the BOLD measurements from cerebrospinal fluid (CSF) and white matter (WM) voxels, as defined by SPM12’s segmentation functionality, were generated using the PhysIO toolbox^85^ and included as nuisance denoising regressors. The PhysIO toolbox was part of the open-source software package TAPAS^86^ (http://www.translationalneuromodeling.org/tapas).

Simultaneously collected physiological data (respiration rate and pulse oximeter readings) were also modeled with the PhysIO toolbox using the RETROICOR^87^ procedure to estimate Fourier expansions of different order for the phases of cardiac pulsation (3rd order, 6 regressors), respiration (4th order, 8 regressors) and cardio-respiratory interactions (1st order, 4 regressors). In addition, PhysIO was used to model respiratory volume per time (RVT, 1 regressor) and heart rate variability (HRV, 1 regressor). In cases when physiological data were not collected for a single run, these twenty regressors were excluded from models. This occurred approximately equally between the two main tasks (24 runs across five participants for the hierarchical task and 25 runs across six participants in the flat task) and infrequently (5% of runs overall).

### Multivoxel Pattern Analyses

#### Decoding Representational Content

To estimate the representational content of the neural activity in each predefined ROI, we implemented a series of cross-validated decoding analyses using the Decoding Toolbox ^88^ in MATLAB. A series of linear support vector machines (cost parameter *c = 1)* were constructed, separately for each task, to test whether specific binary variables (e.g. stimulus features: adult vs child face, indoor vs outdoor scene, low vs high number, etc; response category: circle vs square; motor responses: left vs right key press) were encoded in the set of run-wise multi-voxel beta patterns associated with each trial type estimated using the GLMs described above (GLM1 for task-relevant features and GLM3 for orthogonal, task-irrelevant features). No additional voxel/feature selection was carried out.

We employed linear classifiers under the widely held simplifying assumption that linear kernels approximate a plausible readout mechanism that might be implemented by a single downstream neuron^89,90^. Classification was implemented using a leave-one-run-out cross-validation scheme. Each run was iteratively held out as a test set, while the remaining runs formed the training set used to train the classifier. Therefore, on each cross-validation fold, the classifier was trained on patterns from n-1 runs, and tested on the patterns from the left-out test run. Classifier decoding accuracy was defined as the mean test set accuracy across all cross-validation folds. A cross-validated decoding accuracy greater than chance (50%) provides evidence that the decoded task dimension is represented in the underlying multi-voxel activity.

Decoding accuracies were statistically evaluated at the group level using permutation testing. For each decoding analysis, labels were randomly permuted independently in each functional run to generate 1000 permuted datasets. A classification was then implemented using an identical leave-one-run-out cross-validation scheme on each of the permuted datasets, producing a null distribution of decoding accuracies for every subject, region and analysis. Group-level null distribution was then generated by averaging across subject-specific null distributions. Finally, *p* values were computed as the proportion of decoding accuracies from the group-level null distribution that were lower than the true group-level mean decoding accuracy.

For comparing decoding accuracies across conditions or ROIs, we employed parametric statistical testing using paired *t*-tests or repeated measures ANOVA. All statistical tests were two-tailed. In addition, Bayes Factors (i.e. the ratio of marginal likelihoods) are reported when evaluating evidence in favor or against a null effect. All Bayes Factors are presented as a ratio of the alternate hypothesis over a null hypothesis (i.e. BF_10_). We interpret Bayes factors based on the classification scheme suggested by Lee & Wagenmakers^91^.

For testing whether task-relevant features are encoded, a total of five sets of classifiers were constructed and tested for each participant and task and region, one to decode each of the three stimulus features, one to decode the response category (e.g., square or circle category), and one to decode the motor response (left vs right key press) on the patterns estimated from GLM1. Another set of 3 classifiers was constructed to decode each of the three orthogonal, task-irrelevant stimulus features on the patterns estimated from GLM3.

#### Separability Analysis

To estimate the separability of neural representations in each predefined ROI, we employed a previously described decoding-based approach. Specifically, we examined the decodability of all possible binary dichotomies with a linear classifier. These binary dichotomies reflect the many possible ‘dimensions’ that could be read out from the mixture of the task inputs. The greater the number of such dichotomies that can be read out, the more separable the representation and the higher its dimensionality^34,35^. While a total of 2^8^ possible dichotomies exist given 8 possible inputs, we restricted ourselves to all 35 balanced binary dichotomies which had an equal number of patterns for each class (4 trial types each) ^31,40^ to minimize classifier bias resulting from unbalanced training data. Therefore, for each subject, ROI and task we trained a set of 35 linear classifiers to test the decodability of each of these dichotomies using the same leave-one-run-out cross-validation approach as described above. Mean cross-validated decoding accuracies for each of the 35 linear classifiers were evaluated against chance with a permutation test at the group level after a Bonferroni correction for 35 multiple comparisons.

Separability analyses were separately carried out for each subject, ROI, task, and patterns estimated from GLM1 (trial-types defined by task-relevant features) and GLM3 (pseudo trial-types orthogonal, task-irrelevant features). In addition, task-specific analyses were separately carried out for response-category-specific patterns (flat task) and context-specific patterns (hierarchy task) from GLM2 to estimate the separability of local representations in response-category and context subspaces respectively in the flat and hierarchy task. Given that each of these analyses of local representational geometry relied on data from half the trials as the global analyses, we opted to not apply Bonferroni correction for the 35 multiple comparisons, sacrificing specificity for sensitivity.

We assessed separability using three different measures. First, we computed, at the group level, the proportion of 35 dichotomies could be successfully decoded in a group-level permutation test (after a conservative multiple-comparison correction for all 35 comparisons). Second, we computed *shattering dimensionality*, defined as the mean decoding accuracy across all 35 dichotomies again assessed against chance with a group-level permutation test. Shattering dimensionality reflects the overall separability of the representation and is informed by separability along a wide range of neural dimensions. Because shattering dimensionality can be biased by the strength of pure-selective coding of task variables, it does not cleanly reflect separability resulting from the non-linear mixing of individual input features in the representation. Finally, therefore, to examine separability while minimizing the influence of pure selective coding, we also defined *mixed selective variability* as the mean decoding accuracy across the subset of dichotomies that were orthogonal to all identifiable task-relevant variables (3 stimulus features and response categories). This measure would only be above-chance levels if voxel responses were dependent non-linearly on two or more task-relevant variables and therefore provides a conservative test of non-linear mixing. Separability measures were formally compared across tasks, GLMs or ROIs using two-tailed, parametric statistical tests.

#### Abstraction Analysis

To test if the representation of a task feature was abstract (i.e., invariant to changes in independent task features), we carried out cross-generalization analyses where we tested the classifier on patterns derived from a different set of conditions than on which it was trained, again using a leave-one-run-out cross-validation scheme as before^31^. For example, to test whether the coding of response category (i.e., square or circle category) was abstract in the flat task, we randomly subsampled two of the trial-types associated with each of the two response categories and then tested the classifier on the remaining trial-types. In this example, this was repeated 36 times to cover all possible combinations of train and test sets.

The mean cross-validated performance across all these classifiers is referred to as the cross-classification generalization performance (CCGP) ^31^. Above-chance decoding accuracies, in this case, provide evidence that the representations of the decoded features (e.g., response category) are at least partially invariant to changes in other features (e.g., stimulus features) and rule out a maximally expanded high-dimensional representation. Higher levels of CCGP reflect greater degrees of abstraction in the representation. We evaluated whether CCGP was above-chance levels with permutation testing at the group level as described above.

We assessed CCGP for all task-relevant variables from patterns estimated from GLM1. In addition, task-specific abstraction analyses were separately carried out for response-category-specific patterns (flat task) and context-specific patterns (hierarchy task) from GLM2 to estimate the CCGP of local representations in response-category and context subspaces respectively in the flat and hierarchy task.

#### Representational Alignment Analysis

To test whether two representations are aligned, we first trained classifiers to decode each represented variable and then estimated the angle between their coding axes. The coding axis is mathematically defined by a vector of the weights of a trained linear classifier whose direction is perpendicular to the separating hyperplane learned by the classifier. Two representations are fully aligned if their coding axes are parallel, and they are orthogonal if they are at right angles. Therefore, the angle between the coding axes of a pair of trained classifiers provides an estimate of the degree of alignment of the underlying representations. To estimate the representational alignment, we extracted weight vectors from a pair of trained classifiers from each cross-validation fold and calculated their normalized overlap which equals the cosine of the angle between them.

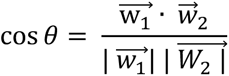

Angles were then computed by taking the arccosine and were averaged across cross-validation folds by computing the circular mean. Angles were statistically evaluated using circular statistics.

#### Representational Similarity Analysis (RSA)

RSA was performed using the python package RSAtoolbox^92^ (version 3.0). First, for each participant and each ROI, a task-level empirical representational dissimilarity matrix (RDM) was constructed by calculating the cross-validated Mahalanobis (‘crossnobis’) distances between the regressors for the eight possible trial types^92^. Cross-validation was performed across run measurements, and the voxel covariance estimates were calculated using the diagonal shrinkage method^93^ on the residual signal from the fit GLM. (see Supplement section S4 for Discussion of the impact of the choice of method for computing voxel covariance estimates).

To test how the representation of different types of task-relevant information might be reflected in the distances between trial-type patterns, we regressed the neural RDMs on a set of separate model RDMs which capture predicted pair-wise distances between patterns driven by three stimulus features and response categories. For the hierarchy task, an additional model RDM was included that reflects the predicted pair-wise distances driven by only the context-relevant feature in each context. All model RDMs are depicted in Fig. S1. All regressions also included an “identity” model RDM that predicts similarity driven only by the individual trial type.

For each participant, we fit the weighted sum of model RDMs which minimizes the cosine distance to their empirical RDM, and the regression estimates are then statistically evaluated using two-tailed, one sample *t*-tests at the group level.

#### Analysis of brain-behavior correlations

To analyze systematic relationship between trial-by-trial variability in neural pattern components in dlPFC and trial-by-trial variability in behavior, we employed linear mixed-effects regression. To obtain trial-by-trial estimates of neural pattern components, we first trained an additional set of support vector machines on single trial estimates obtained from GLM4 to decode task-relevant features using a similar leave-one-run-out cross-validation scheme as described above. In particular, for the flat task classifiers were trained for all three stimulus features and the response category, while for the hierarchy task, classifiers were trained for the context feature, and context-specific classifiers for the relevant and irrelevant stimulus features. For each classifier, single-trial patterns from the left-out test run were then projected onto the feature weight vectors (and bias added) to obtain the signed distance from the classifier boundary. These signed distances (one for each classifier) served as trial-by-trial estimates of the neural components for all the task-relevant variables examined. We then employed mixed effects regression to regress either trial-by-trial, log-transformed response times or choices (using logistic regression) onto neural pattern components. Subject-specific intercepts and slopes were included as random effects. All mixed effects regression was conducted in R with the lme4 library.

## Data and code availability

Preprocessed, deidentified data and analysis code will be made available at the time of final publication on a public repository. Prior to that, data and code are available upon request.

## Author contributions

A.B. and H.K. contributed equally to this work. Project conception: A.B & D.Ba; Methodology: A.B, H.K., D.Ba; Data Collection: H.K.; Analysis and Interpretation: A.B, H.K, D.Bu., D.Ba.; Manuscript writing: A.B, H.K, D.Ba; Funding acquisition: A.B & D.Ba; Supervision: D.Ba.

## Acknowledgements

We thank Stefano Fusi and Matia Rigotti for their helpful advice during the conception of the project, as well as their feedback on methods. We thank the SIEMENS developers for their contribution to the Advanced Physio Logging software which we used to record physiological data. We thank Lynn Fanella, Fabienne McElenry, Michael Worden, Ed Walsh and Jerome Sanes of the MRI Research Facility at Brown University for their expert inputs in MRI protocol design, and cooperation in ensuring smooth data collection. We thank Emily Chicklis, Camillo Aquino, Elizabeth Doss, Sarah Mughal, Tanvi Palsamudram, Roshan Parikh, and Ziqi Zhao for their help with data collection. We thank members of the Badre Lab and the wider neuroscience community at Brown University, particularly Emily Chicklis, Atsushi Kikumoto, Avinash Vaidya, Olga Lositsky, Harrison Ritz, Michael Freund, and Daniel Scott, as well as Brad Postle and Ulrich Mayr for their constructive comments and discussions. This work received funding from the National Institute of Mental Health (R01 MH125497) and the National Institute of Neurological Disorders and Stroke (R21 NS108380) of the NIH, and a Multidisciplinary University Research Initiative (MURI) award from the Office of Naval Research (N00014-23-2792). H.K was supported by an NIH Institutional Training Program for Interactionist Cognitive Neuroscience Grant (T32-MH115895).

## Notes

### Competing Interest Statement

The authors have declared no competing interest.

### Summary of Updates

Results section: Typos corrected. Figure 6 caption: Typos corrected. Abstract: A sentence added to highlight brain-behavior result.

