## Supplementary Material for "Task structure tailors the geometry of neural representations in human lateral prefrontal cortex"

**for**

### S1. Deviations from Scanning Protocol

1. Two participants completed the resting and anatomical scan day on their last session instead of their initial scanning day due to scheduling conflicts.
2. One participant completed all but two scanning sessions within 18 days and then was prevented from completing the final two sessions until an additional month had elapsed due to scanner downtime and scheduling issues. They completed practice blocks as usual and the long break did not appreciably affect their performance.
3. One subject, during a single session, was given practice on the incorrect task; this did not affect their performance inside the scanner.
4. In response to occasional scanner technical issues (5 instances across 5 subjects), participants were not able to complete all five runs in a single session. In these cases, participants would complete six runs in subsequent sessions, but would never complete more than six task runs in a single day.
5. One subject was unable to make up all runs and completed only 24 runs of the flat task.
6. For two subjects, due to a coding error, six and nine runs respectively had identical sequences of ITIs (but unique trial sequences).

**S2. Size of preselected regions of interest (ROIs)**

lPFC parcels were obtained from the Schaefer et al. (2018) parcellation defined in MNI space and were mapped to each subject's native space by applying the inverse of the estimated normalization parameters. Left and right lPFC ROIs used for all analyses were created by pooling all voxels across the left lPFC1-5 parcels and right lPFC1-5 parcels respectively.

| ROI name | Average size (in voxels) | Source |
|---|---|---|
| Left lPFC 1 (parcel no. 128) | 112.95 | Schaefer et al. (2018) parcellation (400 parcel version). Obtained from www.github.com/ThomasYeoLab/Standalone_Schaefer2018_LocalGlobal/ |
| Left lPFC2 (parcel no. 129) | 158.4 | |
| Left lPFC3 (parcel no. 130) | 83.5 | |
| Left lPFC4 (parcel no. 131) | 198.5 | |
| Left lPFC5 (parcel no. 132) | 71.2 | |
| Right lPFC 1 (parcel no. 330) | 83.2 | |
| Right lPFC2 (parcel no. 331) | 84.5 | |
| Right lPFC3 (parcel no. 332) | 100.1 | |
| Right lPFC4 (parcel no. 333) | 106.6 | |
| Right lPFC5 (parcel no. 334) | 96.4 | |
| Left Primary Auditory Cortex | 628 | Neurosynth (www.neurosynth.org) |
| Right Primary Auditory Cortex | 568 | |

**S3. Comparing decoding results on merged lPFC ROIs vs averaging across parcels**

Decoding information from the mid-dorsolateral PFC region that we focus on in this paper has been notoriously hard with fMRI[1], in part because of highly noisy measurements. In order to maximize sensitivity to information contained in lPFC ROIs, we elected to conduct our analyses in merged lPFC ROIs that pooled voxels across 5 lPFC parcels in each hemisphere, especially given that our hypotheses do not distinguish between the different parcels. These parcels all cluster into the same functional network[2]; Control A network identified by Yeo et al. [3] and tend to behave similarly in univariate analyses.

A possible critique of this approach is based on the intuition that information in the brain is organized locally and a readout neuron would likely not receive information from neurons that were being sampled by widely separated voxels. An alternate approach to improve sensitivity, then, is to carry out decoding on patterns estimated from each individual parcel, placing spatial constraints on the features used by the classifiers, and then averaging the decoding accuracies across parcels. To test that our findings are based on local information, we repeated our decoding analyses using this alternate approach.

The results were broadly similar and are reported in the Table below, though we note that the merged ROI approach provided greater sensitivity. This is unsurprising. Decoding accuracies are typically positively correlated with ROI size because, with a larger number of informative voxels, a linear classifier has more evidence to pool. Indeed, the trained linear classifier relies on a principled algorithm for weighting each voxel's evidence. An approach that relies on local information and averaging decoding accuracies involves assigning equal weight to voxels from each parcel rather than letting the classifier optimally weigh and pool the evidence from across parcels optimally. This is the cost one pays for imposing the constraint on 'local' information.

Importantly, the main pattern of results remains largely unaltered as summarized in the table below verifying that our main findings reflect largely local information.

**Decoding Results using a merged ROI and averaged ROI approach**

| | **Flat task** | | | | **Hierarchy task** | | | |
|---|---|---|---|---|---|---|---|---|
| | **Left lPFC** | | **Right lPFC** | | **Left lPFC** | | **Right lPFC** | |
| | Averaged | Merged | Averaged | Merged | Averaged | Merged | Averaged | Merged |
| Visual Feature 1 | 51.8%* | 51.9%* | 50.2% | 52.5%* | 50.6%* | 51.8%* | 50.4% | 50.9% |
| Visual Feature 2 | 51.7%* | 56.2%* | 50.6%* | 51.6% | 50.9%* | 52.7%* | 50.1% | 51.4% |
| Auditory Feature | 51.7%* | 53.1%* | 50.7% | 50.8% | 54.6%* | 61.5%* | 53.9%* | 58.9%* |
| Response category | 51.5%* | 53.9%* | 51.5%* | 52.5%* | 52.2%* | 54.4%* | 51.3%* | 52.2%* |
| Orthogonal VF1 | 49.8% | 50.5% | 50.3% | 50.8% | 49.6% | 49.7% | 50.2% | 50.1% |
| Orthogonal VF2 | 50.8% | 51.3% | 51.1% | 53.6%* | 50.1% | 51.8%* | 50.3% | 50.4% |
| Orthogonal AF | 50.3% | 50.1% | 49.9% | 50.6% | 50.1% | 50.3% | 50.3% | 49.5% |
| Orthogonal RC | 49.5% | 49.9% | 50.0% | 50.8% | 50.0% | 49.7% | 50.1% | 50.5% |
| Separability | 51.0%* | 52.8%* | 50.4%* | 52.6%* | 51.1%* | 51.3%* | 50.7%* | 51.7%* |
| Mixed-selective sep | 50.6%* | 51.5%* | 50.40% | 50.50% | 50.4%* | 50.20% | 50.00% | 49.60% |
| CCGP VF1 | 49.90% | 49.40% | 50.20% | 50.70% | 49.30% | 48.70% | 49.60% | 49.50% |
| CCGP VF2 | 50.40% | 52.1%* | 50.20% | 50.50% | 49.90% | 49.80% | 49.60% | 49.60% |
| CCGP AF | 50.20% | 50.40% | 49.80% | 49.70% | 54.10% | 60.3%* | 52.7%* | 56.90% |
| CCGP RC | 51.2%* | 53.2%* | 51.0%* | 50.80% | 50.30% | 52.4%* | 50.20% | 50.40% |

**refers to a parametric group-level t-test against chance*

**S4. Sensitivity of RSA to noise normalization procedure**

To estimate the Representational Dissimilarity Matrices for RSA it is a recommended practice to pre-whiten the regression estimates that constitute the pattern by accounting for the noise covariance structure across voxels. [4] Such multivariate noise normalization makes the noise component of the patterns approximately i.i.d. However, there is some debate on the pros and cons of multivariate noise normalization given its variable impact on reliability and RSA effect sizes. [4,5] Moreover, there is more than one approach to estimate the covariance structure across the voxels, including relying on residuals of the original GLM used to estimate the patterns or relying on the regression estimates from the GLM themselves. In the manuscript we take the approach recommended by Walther et al. [4] of relying on the residuals (also known as the residuals approach). However, for completion, we note here that we also attempted pre-whitening using a covariance estimate obtained from the regression estimates (also known as the measurements approach).

We note that the main pattern of results remains unaffected by the choice of noise normalization approach, though effects are somewhat weaker when using the measurements approach. In the flat task in the left lPFC, we find unique effects of visual feature 2 ($t = 2.8$, $p = 0.011$*) and the response category ($t = 2.5$, $p = 0.02$*). In the hierarchy task in the left lPFC we find evidence for unique effects of the context/auditory feature ($t = 7.6$, $p < 0.001$*) and weaker effects of response category ($t = 2.25$, $p = 0.036$), and the local context-relevant stimulus feature ($t = 3.6$, $p = 0.002$).

### Supplementary Figures

#### SF1. Model RDMs employed in RSA

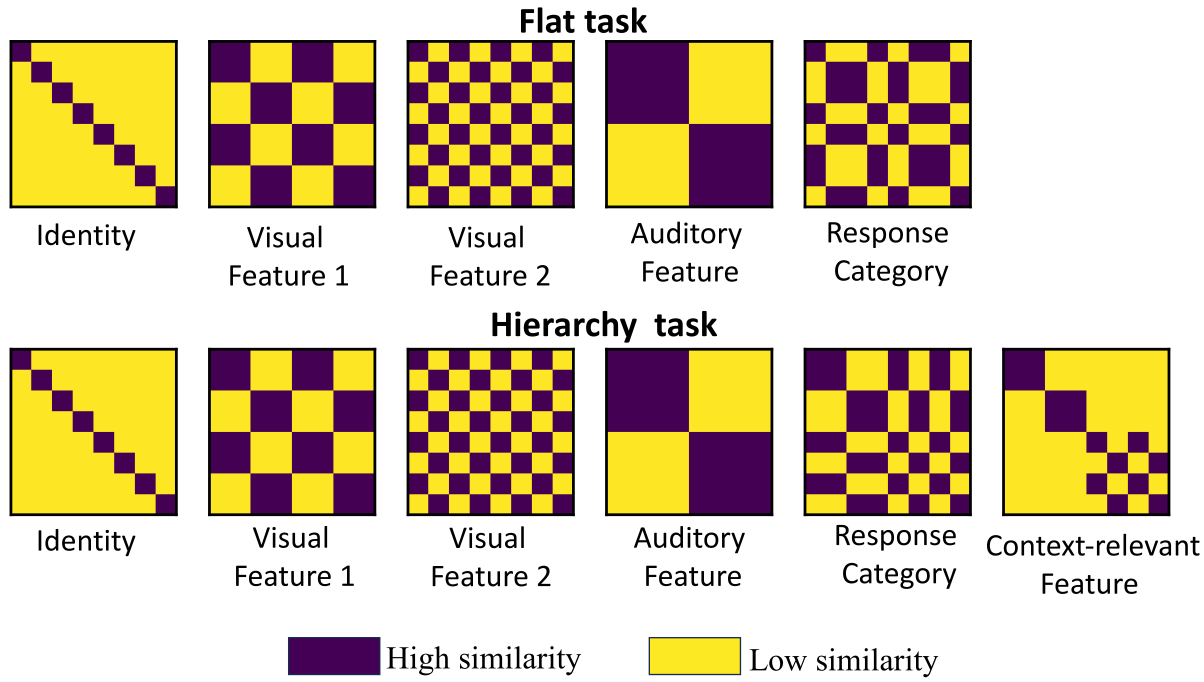

**Figure SF1.** Model Representational Dissimilarity Matrices (RDMs) employed in linear regression analysis of neural representational structure (i.e. neural RDMs). RDMs reflect predicted pair-wise distances between the 8 different trial types. Blue color reflects high similarity (a predicted distance of 0) and the yellow color reflects high dissimilarity (a 'distance' of 1). In the flat task (top row), regressors where dissimilarity was predicted solely by identity (unique trial type), visual feature 1, visual feature 2, auditory feature and response category were included. For the hierarchy task (bottom row), an additional regressor was included where similarity was determined by the context-relevant stimulus feature.

**SF2. Results of Representational Similarity Analyses**

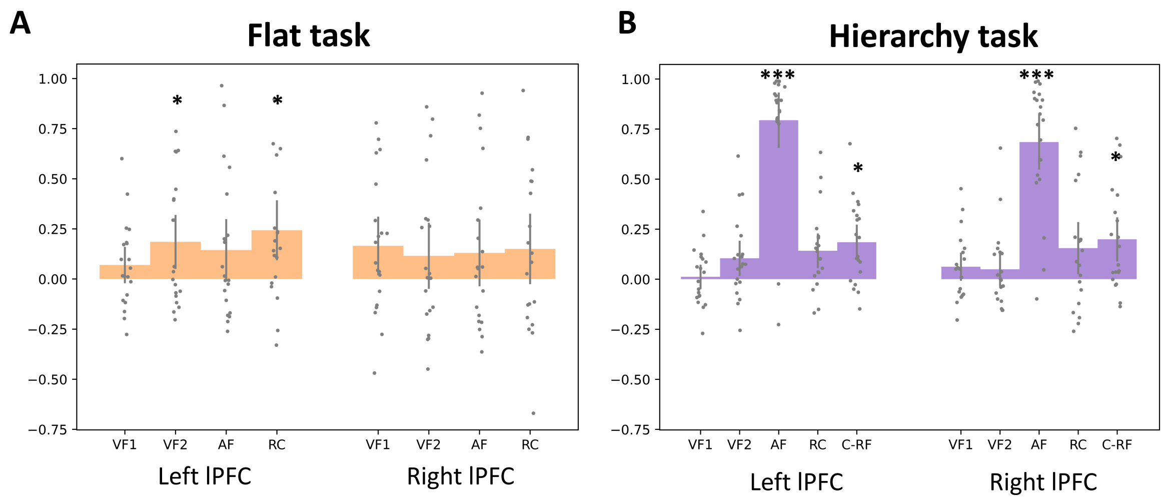

**Figure SF2.** Mean regression estimates from the multiple linear regression of the neural RDM in left and right lPFC on the model RDMs for the flat (A) and hierarchy (B) tasks. Mean estimates reliably different from 0 provide evidence for the unique effect of the regressor. In the flat task, the mean regression estimates for the response category and visual feature 2 were significant in left lPFC. In the hierarchy task, across both left and right lPFC there was a strong effect of the auditory feature (or context) and also the context-relevant stimulus feature (C-RF). Error bars reflect the 95% confidence intervals.

**SF3. Multi-dimensional Scaling of group-level RDMs**

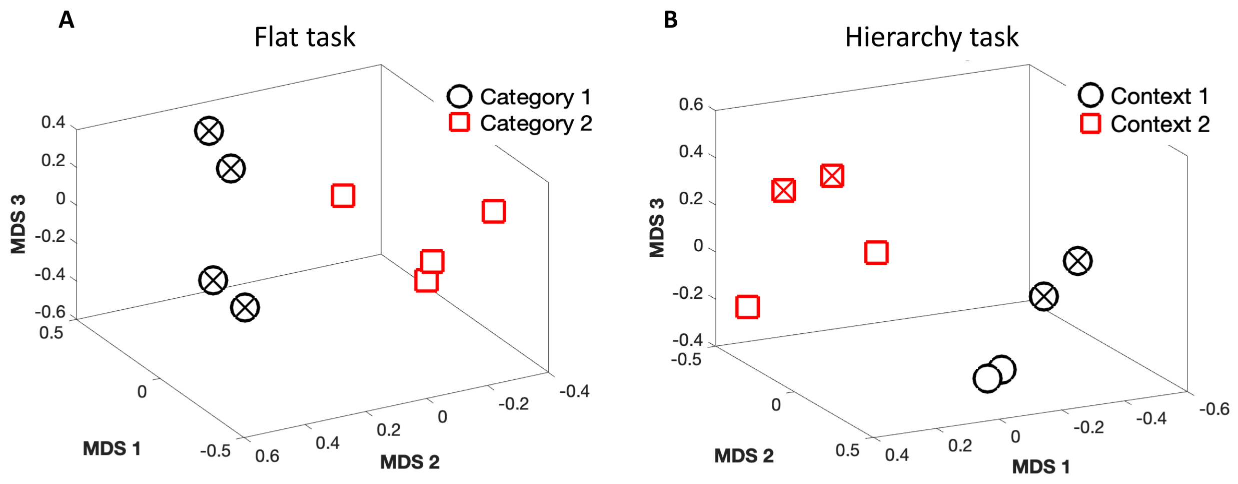

**Figure SF3.** MDS-based visualization of group-mean representational geometries in the flat (A) and hierarchy task (B). In the left lPFC, for each task separately, a group-level representational dissimilarity matrix (RDM) from the subject-specific RDMs used for the RSA. We then employed non-metric multi-dimensional scaling (MDS) to visualize geometries consistent with these RDMs. Non-metric MDS with 3 components, 100 unique initializations (runs), and 10,000 steps per run was computed using the distances in the RDMs. Plots show a 3-dimensional embedding. For the flat task, clustering is by response category and the two response categories are well-separated (red squares vs black circles). For the hierarchy task, clustering is by context (red squares vs black circles), with local geometries reflecting additional clustering by the context-relevant stimulus feature. Markers with and without Xs reflect the response categories across both plots.
